# Decoding mutational hotspots in human disease through the gene modules governing thymic regulatory T cells

**DOI:** 10.1101/2023.12.27.573411

**Authors:** Alexandre A. S. F. Raposo, Pedro Rosmaninho, Susana L. Silva, Susana Paço, Maria E. Brazão, Ana Godinho-Santos, Yumie Tokunaga, Helena Nunes-Cabaço, Ana Serra-Caetano, Afonso R. M. Almeida, Ana E. Sousa

**Author notes:** GlaxoSmithKline Biologicals, SA; Rue de l’Institut 89, 1330 Rixensart, Belgium. Faculdade de Farmácia da Universidade de Lisboa, Av. Professor Gama Pinto 1649-003 Lisboa | Portugal. Laboratory of Animal Genomics, GIGA-Medical Genomics, University of Liege, Belgium.

## Abstract

Computational strategies to extract meaningful biological information from multiomics data are in great demand for effective clinical use. This is most relevant in immune-mediated disorders, where the combined impact of multiple variants is difficult to determine. Regulatory T cells (Tregs), particularly those lineage-committed in the thymus, are essential for immune homeostasis and self-tolerance, controlling inflammatory and autoimmune processes in many diseases with a multigenic basis. Here, we quantify the Transcription Factor (TF) differential occupancy landscape to uncover the Gene Regulatory Modules governing human thymic Tregs, providing a tool to prioritise variants in complex diseases. Combined RNA-seq and ATAC-seq generated a matrix of differential TF binding to genes differentially expressed in Tregs, in contrast to their counterpart conventional CD4 single-positive thymocytes. The gene loci of both established and novel genetic interactions uncovered by the Gene Regulatory Modules were significantly enriched in rare variants carried by patients with common variable immunodeficiency, here used as a model of polygenic-based disease with severe inflammatory and autoimmune manifestations. The Gene Regulatory Modules controlling the Treg signature can, therefore, be a valuable resource for variant classification, and to uncover new therapeutic targets. Overall, we provide a tool to decipher mutational hotspots in individual genomes.

## INTRODUCTION

Immunological and inflammatory diseases are often associated to a complex genetic basis and epigenetic perturbations. Whole-genome sequencing (WGS) has been increasingly used to unravel the multigenic contribution to these disorders. However, the promise of molecular profiling and individual therapies has so far fallen short of expectations (Stessman et al. 2014). Most strategies are based on Genome-Wide Association Studies (GWAS) and many rare Single-Nucleotide Variants (SNV) found are of unknown significance (VUS) or correspond to gain-of-function variants not addressed by annotation. Also, SNVs falling in non-coding regions are often insufficiently indicative of causality (Li et al. 2010). Finally, and most importantly, modelling of combined impact of multiple SNV is challenging, with latest research limited to digenic systems (Kerner et al. 2020). Integration of WGS with gene regulatory networks addresses both issues by aggregating weak genetic signals through independent evidence of causal link (Zhu et al. 2021). Such strategy may uncover previously undescribed trait-associated interactions and provide a way to prioritise variants and reveal therapeutical insights. This is of utmost importance given the wide variety of phenotypes associated with many immunological and inflammatory diseases.

CD4 T cells are the main organisers of immune responses. They are essential to mount effective antibody responses, to promote the generation of cytotoxic lymphocytes targeting tumors and infected cells, and to govern the innate immune responses (Nunes-Cabaço et al. 2011). Therefore, CD4 T-cell disturbances are likely to have a crucial impact in the outcome of immune disorders. CD4 T cells are divided in effector conventional (Tconvs) and the suppressive regulatory T cells (Tregs). They develop primarily in the thymus, although Tregs can also be induced from Tconvs in the periphery *(Silva and Sousa 2016; Silva et al. 2016)*. Thymic Tregs (tTregs) are believed to be enriched in self-reactive TCRs, which is thought to further enable them to limit auto-reactive responses, and, therefore, are particularly relevant for self-tolerance and immune homeostasis (Caramalho et al. 2015a). Identifying the regulatory modules that control the Treg signature in the human thymus is crucial to reveal factors whose deregulation may play a role in immune pathology (Mijnheer et al. 2021). Despite this, the focus has been so far on peripheral Tregs, including both thymic-derived and peripherally-induced Tregs (Mijnheer et al. 2021; Ohkura and Sakaguchi 2020). Moreover, such studies fail to explore the chromatin accessibility landscape of tTregs (Mijnheer et al. 2021; Rodriguez et al. 2015). In the thymus, single-cell sequencing has been employed in the characterisation of early T-cell commitment and organogenesis, both in mice and humans (Morgana et al. 2022; Cordes et al. 2022; Giladi et al. 2018; Zhou et al. 2019; Zeng et al. 2019; Park et al. 2020). Although this technique allows the profiling of heterogeneous, rare cell populations, and their developmental dynamics, it cannot yield the sequencing depth achieved by bulk RNA-seq, and does not warrant full coverage of the universe of transcripts (Van Der Wijst et al. 2018), nor the sensitivity required by second order analyses of ATAC-seq data such as TF occupancy (Bentsen et al. 2020). Altogether, there is a need for genome-wide data on chromatin accessibility in human tTregs, and higher resolution analyses, supported by quantification methods, to understand how tTreg-specific gene expression is regulated. Lineage-specific expression is regulated by lineage-enriched binding of multiple TF (Transcription Factor Binding Sites, TFBS) to cis-regulatory elements in the genome (Rothenberg 2021; Little et al. 2021). ChIP-seq is the most established technique to fully map and quantify TFBS. Still, it is demanding in cell numbers. Alternative techniques, such as CUT&RUN (Skene and Henikoff 2017), also require TF-specific antibodies which limit studies to a few tens of TF regulators. ATAC-seq provides a comprehensive alternative: combined with appropriate digital genomic footprinting of regions of open chromatin (ROC), it allows the compilation of the full cis-regulatory elements, as well as an estimate of TF occupancy at the respective genomic site (Bentsen et al. 2020; Buenrostro et al. 2015). It is assumed that TF occupancy/binding is a measure of TF activity and differences in TF binding at the same TFBS can be informative on the activity of the same TF in different lineages (Little et al. 2021). This quantification, however, is so far surprisingly absent in published regulatory models for gene expression during thymocyte development (Shin and Rothenberg 2023).

Here we defined the expression signature that distinguishes tTregs from tTconvs and quantified genome-wide TF binding. Applying an artificial intelligence approach to TF differential binding maps, we uncovered the main Gene Regulatory Modules (GRM) shaping the identity of tTregs in the human thymus. We tested whether these GRM generated from healthy thymuses are predictive of mutational hotspots in a cohort of patients with Comon Variable Immunodeficiency (CVID), here used as a model for complex immune diseases (Lopes-da-Silva and Rizzo 2008; Silva et al. 2019; Berbers et al. 2021). The whole genome sequencing (WGS) datasets of patients with a disease with a likely polygenic basis provide unique testing data to validate the model by inferring the combined impact of rare SNV. Additionally, the integration of WGS data also allowed us to ascertain the biological relevance of the GRM model. Thus, the GRM of human thymic Tregs delivers both a blueprint for the genome-wide transcriptional programme defining the Treg lineage as well as a tool to help categorise multiple rare variants in immune disorders.

## RESULTS

### The human thymic Treg signature and landscape of accessible chromatin

Regulatory T cells, particularly those committed in the thymus, play a non-redundant role in the control of autoimmune and inflammatory diseases. Therefore, it is critical to identify the relevant networks of epigenomic interactions governing thymic Tregs (tTregs), or “Gene Regulatory Modules” (GRM). For this purpose, we used the genome-wide expression (RNA-seq) and chromatin accessibility maps (ATAC-seq) of purified CD4 single-positive (CD4SP) Treg and Tconv cells from human thymuses (Fig. S1A). RNA-seq yielded ca. 13,000 genes with non-neglectable expression levels in at least one of the lineages (E-MTAB-11211, Fig. S1B), whilst peak-calling of ATAC-seq signal identified 188,169 Regions of Open Chromatin (ROC, E-MTAB-11220, Fig. S1C), including in promoters, gene body, or intergenic regions (Raposo et al. 2015).

We first investigated the association between the 1,357 Differentially Expressed Genes (DEG) defining the “tTreg Signature” (Supplemental Table S1) and the ROC to model the regulatory topology controlling it at the transcriptional level. Quantification of differential chromatin accessibility (DCA) between the tTreg and tTconv resulted in 90,437 regions more open in tTregs (“Open” ROC, DCA > 0); and 97,732 more open in tTconv (“Closed” ROC, DCA < 0) (Fig. 1A; Fig. S1C; Supplemental Table S2). Based on annotation to the nearest TSS, 8,062 of these ROC were potentially regulating differential gene expression, with a median of 4 ROC per gene (Supplemental Table S2, ROC annotation/DEG overlap of 1,265/1,357=93%). Since ROC bearing both significant and non-significant differences in accessibility may influence differential expression, all those associated to DEG were included in the remaining analysis and defined as the “tTreg chromatin landscape” (Supplemental Table S2).

**Figure 1.**
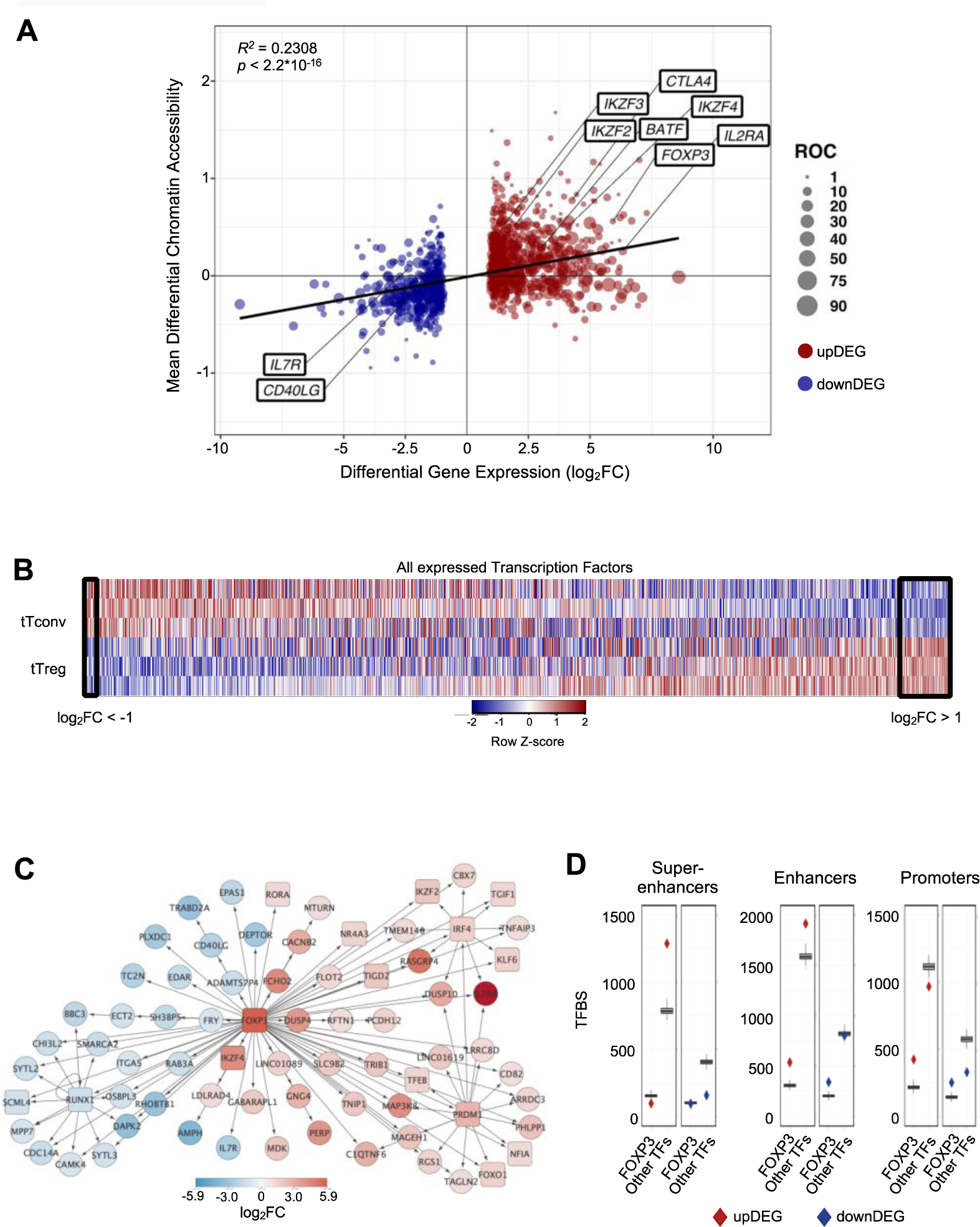
The human thymic Treg signature, correlation with chromatin accessibility and transcription factor binding. **A.** Correlation between Differential Gene Expression (DGE, FDR<0.05, |log_2_FC|>1, RNA-seq) and Differential Chromatin Accessibility (DCA, q-value < 0.05, ATAC-seq) between regulatory (tTreg) and conventional (tTconv) thymocyte; black line, linear regression, *R^2^*=0.2308, *p*<2.2*10^-16^; bubbles on mean DCA (log_2_FC) for all the ROC associated to each Differentially Expressed Gene (DEG, log_2_FC), with size of bubble proportional to number of regions; labels refer to known markers of either lineage. **B.** Expression levels for all transcription factors (TF) in tTreg and tTconv data sets (TF, n=1,010, log_2_CPM shown in column Z-score); left box referring to 16 TF down-regulated in tTregs (downDEG) and right box indicates 56 TF up-regulated in tTregs (upDEG), with *FOXP3* as the TF with highest fold-change. **C.** FOXP3 regulatory network: arrows represent binding of TF (square nodes) to their target DEG (circular nodes). **D.** Frequency of Transcription Factor Binding Sites (TFBS) mapping to Treg-specific super-enhancer, enhancer, and promoter regions, associated to FOXP3 targets (FOXP3 and co-binding), or other targets; boxplots refer to 1.5xIQR. Data refer to 3 replicates generated from 3 different thymuses. Blue - downDEG; red - upDEG (except in B). See also Fig. S1 and S2, and Datasets 1-4.

Open/closed ROC directly correlated to up/down DEG (from now onwards, “upDEG” and “downDEG”), respectively, as shown by regression analysis of DCA vs DEG (*R^2^* = 0.2308, *p* < 2.2*10^-16^, Fig. 1A and Supplemental Table S2), showing that differential chromatin accessibility may be predictive of differential gene expression.

However, chromatin accessibility of many of the annotated ROC (73% with | log_2_DCA | < 0.5, from a total of 3,593 open + 3,927 closed) varies little between lineages, suggesting further layers of transcriptional control localising to these regulatory sites.

The tTreg Signature (Supplemental Table S1) includes 56 up-regulated and 16 down-regulated transcription factors (TFs), with *FOXP3*, the master Treg TF, as the most upregulated (log_2_FC=5.93, Fig. 1B). In addition, tTregs and tTconvs express many other TFs (Fig. 1B), which may also contribute to define the Treg identity. We asked which TFs are targeting tTreg signature genes using digital genomic footprinting of ATAC signal, both at tTreg and tTconv ROC. The corresponding TFs were then identified by crossing the JASPAR database of consensus sequences for 639 TFs against the 1,010 TFs that are expressed in human CD4SP thymocytes (Li and Altman 2018). We thus identified 34,167 TF binding sites (TFBS) in the tTreg chromatin landscape bound by one of the 233 TFs expressed in tTregs, with each TF-TFBS interaction estimated by its occupancy (TFBS binding score, Fig. S2A, and Supplemental Table S3).

We found that FOXP3 binds directly to 87 TFBS in tTregs, potentially regulating 74 targets (Fig. 1C, Supplemental Table S3 and S4). From these, 44 are upDEG and include known Treg markers, and many currently unreported transcripts potentially required for Treg identity in the human thymus. Conversely, we found *IL7R* and *CD40LG* amongst the 30 downDEG directly bound by FOXP3 (Fig. 1C, Supplemental Table S4). FOXP3 downstream direct regulation includes potential FOXP3 binding to TFs which may have their own direct targets downstream of FOXP3 (*RORA*, *IKZF4*, *NR4A3*, *TGID2*); and to three subnetworks of co-regulation (Fig. 1C, Supplemental Table S4), defined as simultaneous FOXP3 binding to a TF and its targets (Trujillo-Ochoa et al. 2023). Specifically, we found FOXP3 TFBS at the repressor *RUNX1,* and at their co-downregulated targets (Ono et al. 2007); at *IRF4* and their co-regulated genes, including *IL2RA* and *IKZF2*; and other TFs with no DEG targets in common with FOXP3, such as *KLF6* and *TGIF1.* Finally, FOXP3 binds to up-regulated *PRDM1*, which protein our data predicts to co-bind several FOXP3 direct targets, including *FOXO1* – which might constitute a forward feedback loop for FOXP3 (Kerdiles et al. 2010; Trujillo-Ochoa et al. 2023) (Fig. 1C, Supplemental Table S4).

We found that FOXP3 targets are rarely bound by TFs at their super-enhancers in Tregs, regardless of direction of regulation (Fig. 1D). This is interesting given the relevance of super-enhancers, H3K27ac-rich regions densely bound by master TFs (Hnisz et al. 2013), in defining lineage commitment (Kitagawa et al. 2017). In contrast, we found that TFBSs associated to upregulated DEGs not bound by FOXP3 (“other targets”) are mapping significantly to super-enhancers and enhancers in Tregs (Fig. 1D, Figs. S2B and S2C). Thus, regarding the epigenetic regulation of the human tTreg expression signature, we found FOXP3 binding to the promotors of a small fraction of the targets, and that the upregulation of tTreg genes with no FOXP3 direct binding vastly relies on the occupancy of tTreg-specific super-enhancers by many other TFs.

Overall, these results indicate that the regulation of transcription for tTreg Signature genes may depend, in part, on increased/decreased accessibility in tTregs to specific ROC. Our analysis of binding patterns uncovered a TF transcriptional program distinct from that of FOXP3 and is suggestive of another layer of regulation for the tTreg expression signature(Trujillo-Ochoa et al. 2023).

### TF Differential Binding reveals main Gene Regulatory Modules controlling the tTreg Signature

Next, we reasoned that assessing differential binding associated to DEG may uncover the most relevant modules of the tTreg gene regulatory network. We scored the tTreg vs tTconv differential binding for the 233 candidate TFs (Fig. S2A), at each of the 22,180 and 11,987 TFBS detected in association with the upDEG and downDEG, respectively (Supplemental Table S3). To determine the TF groups associated with different chromatin landscapes potentially regulating the tTreg signature and their target DEG, we considered only sites with a significant binding score in tTregs (ie, “TFBS tTreg bound”, *p* < 0.01) and used *k*-means for unsupervised double clustering of TF and DEG according to their TF differential binding profiles (Figs. S3 and S4; Supplemental Tables S3 and S5). This strategy allowed us to identify the Gene Regulatory Modules (GRM) – defined as the pairing between a TF cluster and a DEG cluster – with the highest differential binding densities in upDEG and downDEG (Fig. 2). The top GRM comprise: the AP-1 family; the ETS domain family; and the KLF/SP protein family (Fig. 2; Figs. S3-S5; Supplemental Table S5).

**Figure 2.**
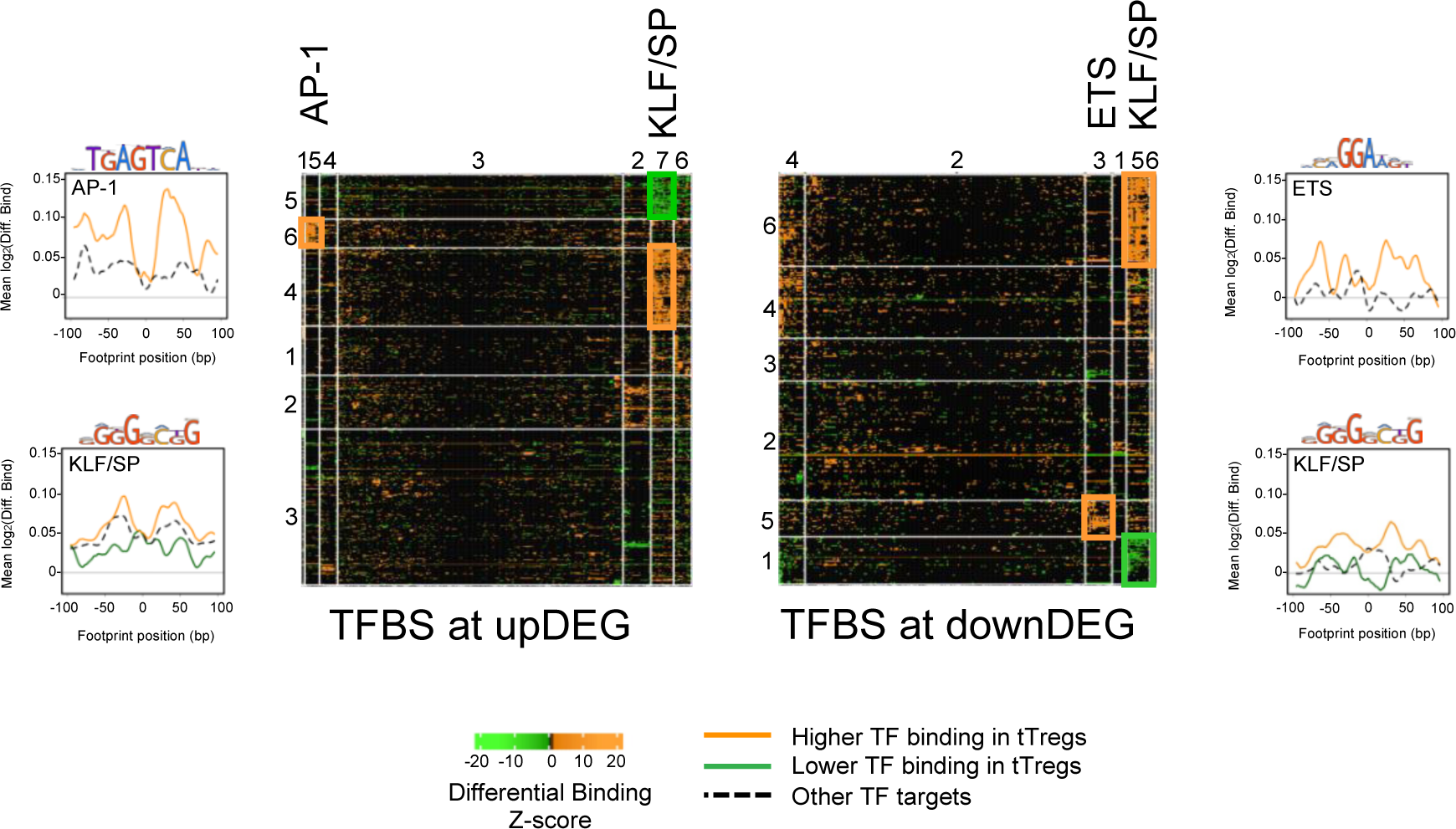
Transcriptional Factor Differential Binding reveals main Gene Regulatory Modules controlling the human thymic Treg Signature. Heatmaps of differential binding score of expressed Transcription Factors (TF) bound to Differential Expressed Genes (DEG), upDEG (top) and downDEG (bottom); *k*-means double clustering, Transcription Factor Binding Sites (TFBS) in columns, respective bound targets, or DEG, in rows; the significant Gene Regulatory Modules (GRM) are highlighted with the name of the TF cluster on the top of the column; heatmap cells show the mean score for all binding sites of each TF to each DEG. Side graphs show the representative consensus motif for TF family and the profiles of mean differential binding to respective DEG from the GRM within 200bp centred at the respective footprint (LOESS curves), with the black and dashed line representing the mean differential binding of all other DEG clusters targeted by the same TF cluster (background for upDEG in the top and downDEG in the bottom). All panels and graphs: orange, increased binding in tTregs; green, decreased binding in tTregs. See also Figs. S3-S5, and Dataset 8.

The AP-1 TFBS cluster (Fig. 3 and Supplemental Table S5) is formed by BATF (log_2_FC=3.01), MAFK, BACH2, FOSL2, FOS, JUNB, and JUND, which featured high differential binding in tTregs to a cluster of 35 upDEG that included the Treg lineage marker *CTLA4* (Walker 2013); the cytokine receptors *IL15RA* (Caramalho et al. 2015b)*, IFNLR1* and *IL4R*; *PRDM1*; *RORA*; genes coding for proteins involved in cell trafficking, such as *PERP, CDH1, PCDH12*; and the chromatin remodeller *HDAC9*.

**Fig. 3.**
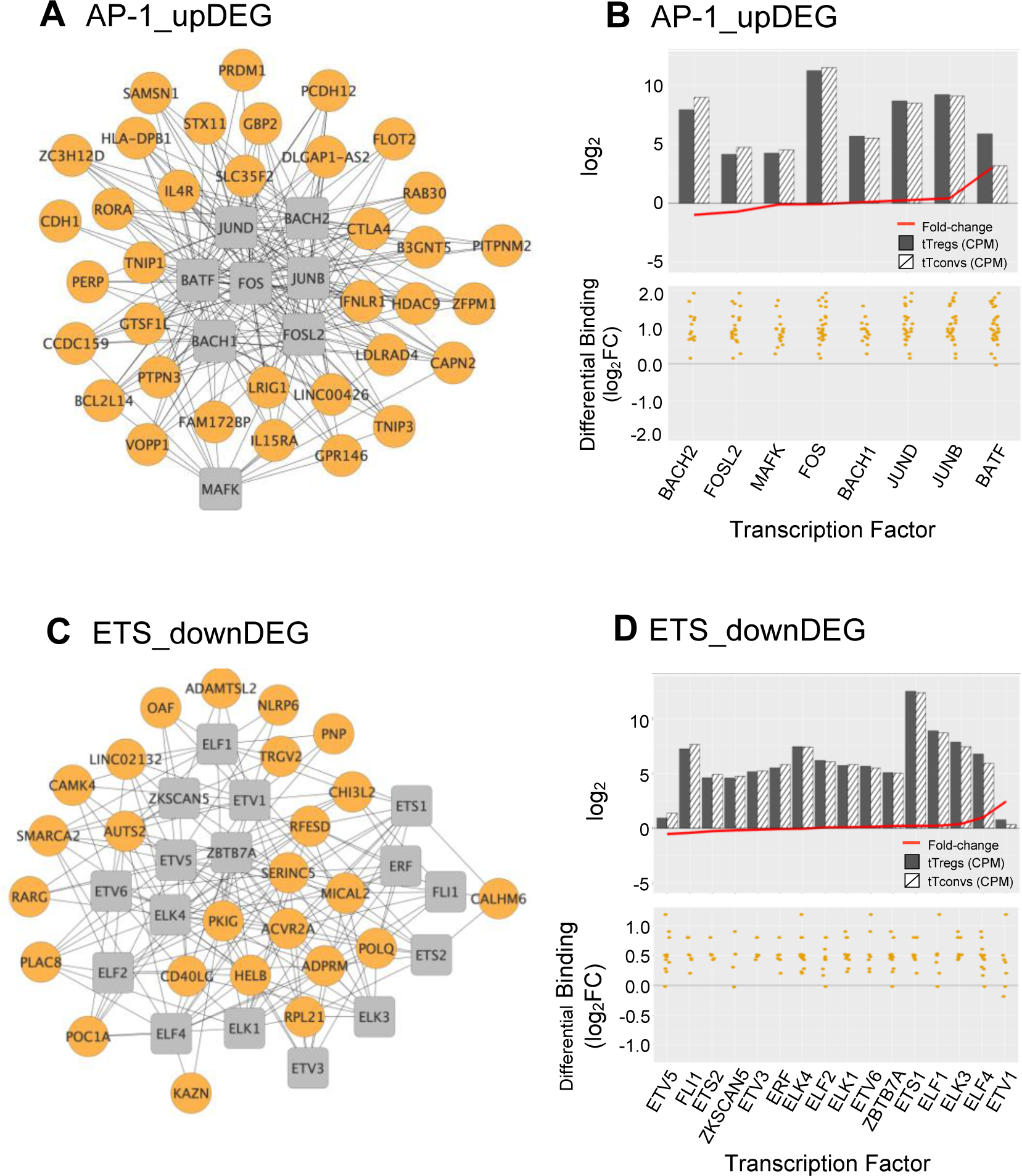
AP-1 and ETS Gene Regulatory Modules and respective Transcription Factor Differential Expression in human thymic Treg signature. A and C. Interactomes of the AP-1 (A) and ETS (C) Gene Regulatory Modules (GRM) representing direct binding by the Transcription Factors (TF) in the respective cluster to their target genes; TF nodes, grey squares; non-TF nodes, orange circles, higher binding by respective TF in tTregs. **B and D**. Analysis of the average expression of the TF included in the AP1 GRM (B) and ETS GRM (D), as well as their differential binding to respective targets. The barplots in the top graphs show TF Average Expression (log_2_CPM) in tTreg (black) and tTconv (hatch), superimposed to TF Differential Expression (red lineplot, log_2_FC); the bottom graphs show the Transcription Factor Differential Binding in tTregs to targets by TF in these GRM.

The ETS domain TF cluster is characterised by high differential binding in tTregs in downDEG (Fig. 3C, Supplemental Table S5), and includes ELF2, ETS1 and 2; ETV5 and 6, ELF1, ELK1, ELK3 and 4, FLI1, chromatin remodeller ZBTB7A (or LRF, partner to ZBTB7B/Thpok) (Yu et al. 2020); ZKSCAN5; and ETV1 and ELF4, significantly upregulated in tTregs (log_2_FC=2.44 and log_2_FC=1.01, Fig. 3D and Supplemental Table S5). Notably, the ETS cluster binds directly to the Tconv lineage marker *CD40LG* (Elgueta et al. 2009); and to *RARG*, which binds to the Foxp3-CNS1 to maintain peripheral Tregs (Maruyama et al. 2011).

The third TF cluster is composed of KLF/SP family, which may act as transcriptional activators or repressors (Presnell et al. 2015; McConnell and Yang 2010). The transcriptional activator KLF6 is the only TF significantly upregulated (log_2_FC=1.45), although this cluster includes several other TF also expressed in tTregs (Fig. 4A and Supplemental Table S5). The KLF/SP cluster forms distinct GRM, according to higher or lower differential binding to corresponding targets in four clusters of differential expression – two upDEG (Fig. 4B and Supplemental Table S5) and two downDEG (Fig. 4B and Supplemental Table S5). The KLF/SP cluster targets with increased differential binding in tTregs the upDEG cluster which includes *BCL3*, *DUSP4, IL10RA,* the NF-#x03BA;B pathway member *RELB*; and *NFKBIZ,* amongst other NFKB2 pathway inhibitors; and the adhesion molecule *CAV1* (Fig. 4B). The same TF cluster also targets a second cluster of upDEG,

**Figure 4.**
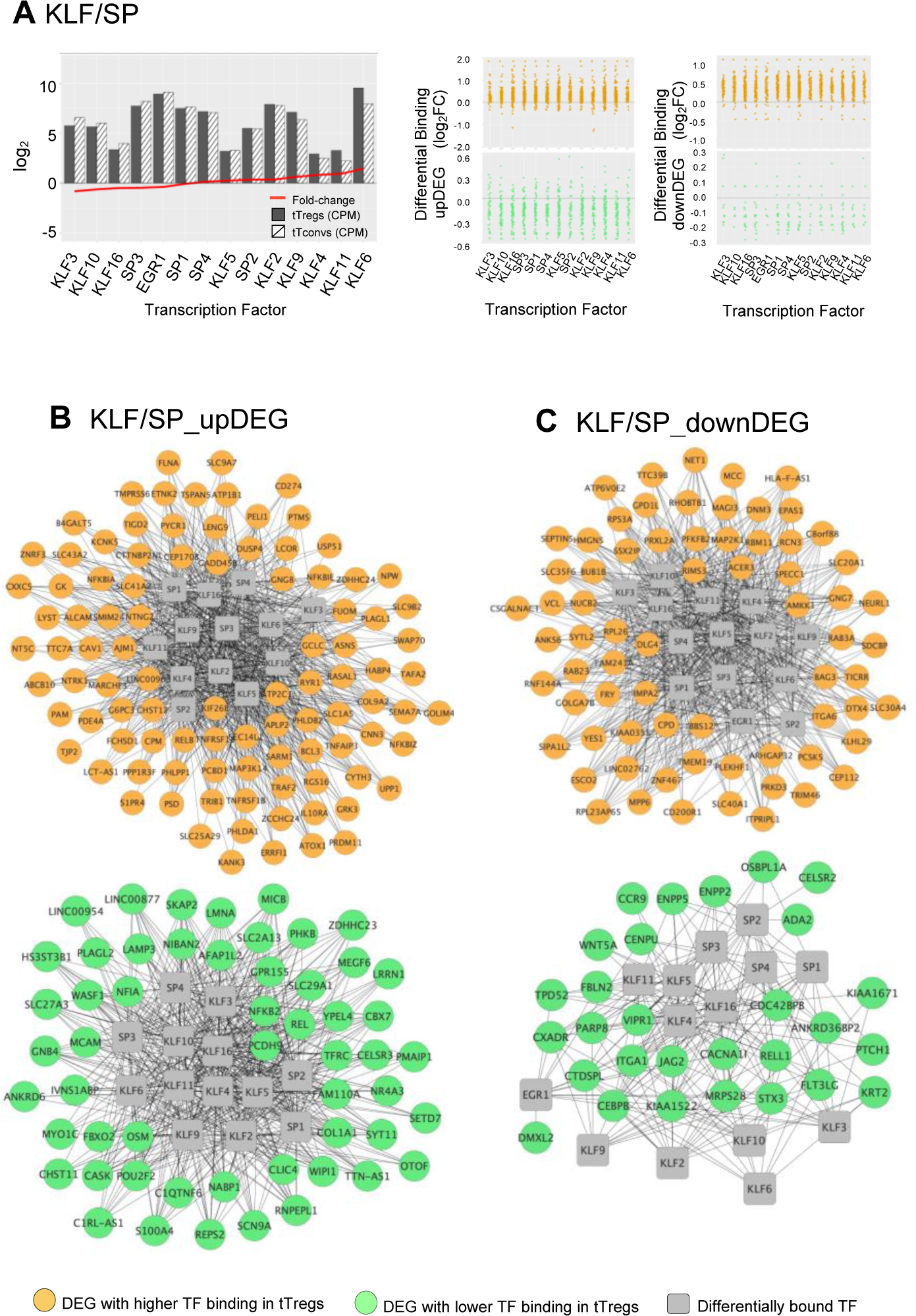
KLF/SP Gene Regulatory Modules and respective Transcription Factor Differential Expression in human thymic Treg signature. **A**. The barplot in the left graph shows TF Average Expression (log_2_CPM) in tTreg (black) and tTconv (hatch), superimposed to TF Differential Expression (red lineplot, log_2_FC); Differential Binding (jitter plots) to targets by TF in the KLF/SP GRM; higher differential binding (orange) in tTregs and lower differential binding in tTregs (green) in upDEG (centre graphs) and downDEG (right graphs). **B.** Interactome of the KLF/SP GRM in upDEG (left) and downDEG (right), representing direct binding by TF in the KLF/SP cluster and respective target genes with higher differential binding in tTregs (orange) and lower differential binding in tTregs (green); TF nodes, grey squares; non-TF nodes, circles in orange for higher binding and green for lower binding by respective TF in tTregs. but with decreased binding in tTregs (Fig. 4B), including *NFKB2, REL*, both NFKB2 pathway members and *POU2F2 (Oct2)*; chromatin organiser *LMNA*; *NR4A3*, a transactivator of *FOXP3* expression; the procadherin *PCDH9*; and *NFIA*, a putative pioneer factor.

Amongst the downregulatory modules (Fig. 4B), high differential binding of transcriptional repressors in tTregs (eg, *KLF9* and *KLF11*) may explain the differential expression of a cluster of 70 downDEG (Fig. 4B), *e.g., DNM3*, a minus-end oriented microtubule molecular motor; integrin *ITGA6*; *EPAS1,* a bHLH factor indispensable to Treg function in mice (Hsu et al. 2020); and *CAMKK1*. Conversely, decreased binding of transcriptional activators (eg, *KLF6* and *KLF3*) in tTregs is a potential mechanism of regulation of 32 downDEG (Fig. 4B), including *CCR9*; *ITGA1*; *WNT5A*; *CXADR;* and *CEBPB*, involved in Tconv differentiation.

We defined the GRM based on direction of expression of targets and the differences in TF occupancy between lineages, which are likely to be further modulated by the TF expression levels, pointing to the TF with highest expression in tTregs, namely *BATF*, *ETV1*, and *KLF6*.

Altogether, taking profit of TF differential binding to define gene regulatory networks more accurately, as it allows us to model DNA-protein interaction directly from genome-wide quantification at a local level, we were able to identify six Gene Regulatory Modules governing the human thymic Treg signature.

### Exploring the Regulatory Networks of human thymic Tregs to decipher complex immune disorders

Given the importance of tTregs in immune-based disorders, the identification of their main regulatory modules is likely to provide a functional tool to prioritise variants when a multigenic cause is expected. To test this possibility, we selected patients with Combined Variable Immunodeficiency (CVID), the most frequent symptomatic primary immunodeficiency (PID). No monogenic cause has been identified in 75 to 95% of CVID cases (Bonilla et al. 2016), suggesting a polygenic basis, as illustrated by our own study in monozygotic twins (Silva et al. 2019). Although the main diagnostic criteria are based on impaired antibody production, severe immune-dysregulatory and inflammatory manifestations featured by CVID patients may be driven by T-cell defects (Motta-Raymundo et al. 2022; Silva et al. 2019; Berbers et al. 2021). Therefore, the tTreg GRM offer a strategy to infer biological meaning from the SNVs, including non-coding mutations documented in CVID patients. To evaluate this possibility, we explored the mutational landscape obtained by whole-genome sequencing (WGS) of 35 CVID patients featuring severe clinical inflammatory/autoimmune phenotypes (Supplemental Table S6).

We hypothesised that tTreg GRM are particularly enriched in rare variants (Pickrell 2014). We focused on rare SNVs (non-Finnish European allelic frequency, AF_NFE < 0.01) and excluded indels and larger structural variations from this analysis. We quantified both the fraction of genes carrying at least one SNV and the number of SNVs per 100kb, since the mutation load estimative is a common approach to evaluate the weight of disease-associated variants in a panel of genes (Sun et al. 2022). Numbers obtained in the GRM associated genes were compared to those obtained with the total tTreg signature genes and the universe of expression in our mature CD4SP thymocyte datasets (see Supplemental Table S7). Results were further compared with those from *de novo* haplotype calling on WGS from blood cells of 35 healthy individuals of Iberian background (WGS data from IGSR, see Supplemental Table S8).

We found that the fraction of genes with at least one rare SNV in CVID patients was significantly higher for GRM genes, performing better than the other gene sets considered (tTreg Signature and All Genes) in capturing the extent of core genes possibly affected in CVID (Fig. 5A, all comparisons p<10^-5^). Importantly, and although a similar result is obtained for healthy individuals, the SNV-gene fraction was always higher in CVID patients when compared to the HC cohort (all p=1.9*10^-12^). In addition, we determined for each of the GRM gene the percentage of CVID and HC individuals featuring at least one SNV, and found, for the large majority of GRM genes, a higher prevalence of mutations in the CVID cohort (Fig. S6, Supplemental Table S9). Mutated genes in the CVID cohort were overrepresented in each of the distinct GRM, both for upDEG and downDEG (Fig. 5B, all comparisons p<10^-10^, see Supplemental Table S7), and there were patients harbouring rare mutations in more than 60% of genes in some GRM (*eg*, KLF^high^_downDEG – Supplemental Table S7). We then questioned if there were specific GRMs more affected by SNVs in some patients than others. To do this, we grouped the CVID patients via hierarchical clustering analysis of prevalence of mutated genes in each GRM (Fig. 5C and Supplemental Table S7). The main difference seems to be established between those patients with an enrichment for the AP1_upDEG GRM and those without (first branching, Fig. 5C), although we could ultimately distinguish 6 clusters of patients (Fig. 5C).

**Figure 5.**
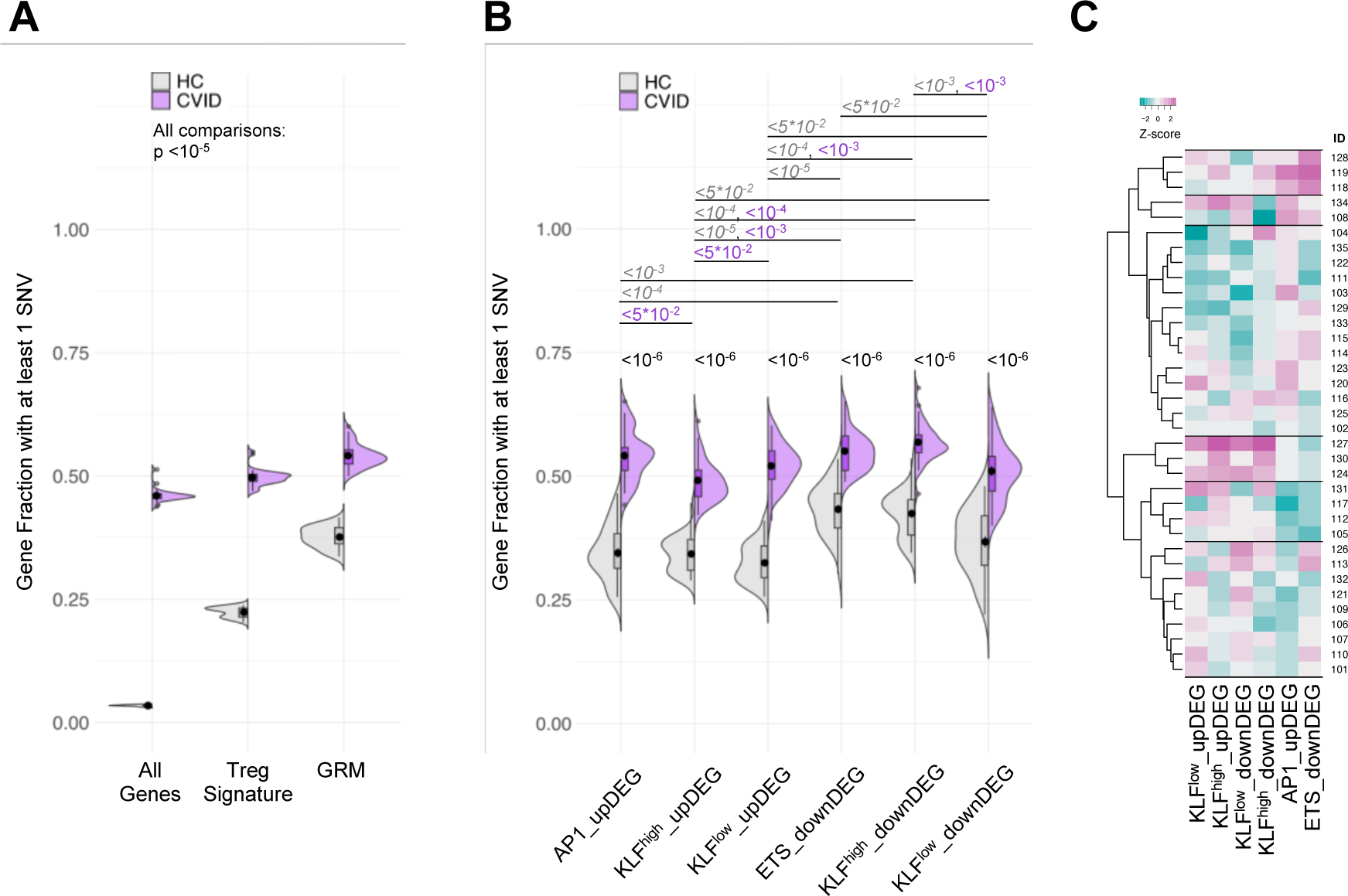
Gene Regulatory Modules of tTreg are enriched in rare variants in CVID patients. **A.** Fraction of genes with at least one SNV in their loci; comparison between distributions for the healthy control cohort (HC, grey) and CVID cohort (purple), in: all genes expressed in tTregs and tTconvs; genes of the tTreg Signature (DEG); and genes forming the GRM. **B**. Fraction of genes with at least one SNV in their loci for each of the GRM (identity indicated), compared amongst them and between CVID and HC cohorts. **C**. Hierarchical clustering of CVID patients by fraction of genes with SNVs in each GRM; six clusters were identified, with the most distinctive patterns set by high prevalence of variant-burden genes in AP1_upDEG and ETS_downDEG (1^st^ cluster from top) and in KLF GRM (4^th^ cluster from top); scale, Z-score: darker magenta, higher fraction; darker cyan, lower fraction. SNV were defined as rare variants (gnomADg, AF_NFE< 0.01) excluding synonymous nucleotide polymorphisms. HC genomes obtained from sample collection of Iberian individuals (IGSR).

Additionally, we estimated the mutation load and made the same analysis, which produced concordant results (Fig. S7). The CVID cohort featured much higher values of SNV counts per 100kb than healthy controls in all gene sets (p=2.0*10^-^12, Fig. S7A). All GRM contributed to this high variant density in CVID (p<10^-5^, Fig. S7B). When clustering CVID patients according to the mutation load in each GRM, we observed two clusters and, again, with a main segregation imposed by the AP1_upDEG GRM (Fig. S7C).

Finally, and given the importance of TFBS in the definition of the GRM, it would be reasonable to assume that these are sites of significant accumulation of variants in CVID patients. Surprisingly, we only found 3 SNVs falling on such TFBS: rs535861886 in patient 103; rs74639548 in patient 113; and rs536121979 in patient 109. These affect binding sites occupied by EGR2 (103 and 113) and SP1 (possibly SP2, 3, or 4, patient 109).

These results provide experimental support to the GRM as a reliable methodology for the integration of T-cell multiomics. The tTreg GRM are significantly enriched in genic regions overlapping rare SNVs found in CVID patients, strongly suggesting that these Gene Regulatory Modules and downstream pathways, which are associated to tTreg identity and function, are disrupted in patients with CVID. These data support the use of differential TF binding and GRM as a tool to assist in the investigation of core genes or pathways underlying the pathogenesis of complex immune disorders.

## DISCUSSION

The definition of GRM by unsupervised clustering of differential binding provides a quantified approach to identify the transcriptional program controlling the tTreg lineage. We uncoupled cellular TF expression from localised binding to the genome, allowing a direct correlation between TF differential binding and target differential expression. The identified modules included genes with recognised prominent role in Treg function and are enriched in variants in CVID patients with clinical evidence of immune dysregulation. We thus propose differential binding as a *bona fide* measurement of TF specific activity, which can overcome limitations in traditional computational inference approaches to regulatory networks.

Modelling of Gene Regulatory Networks via bulk or single-cell ATAC is usually based on a few TF, selected by the top frequency of consensus motifs in regions of open chromatin, enrichment at promoters, or expression level of respective TF (Chopp et al. 2020; DiSpirito et al. 2018; Miraldi et al. 2019; Bravo González-Blas et al. 2020; Shin and Rothenberg 2023). Our novel approach, based on differential TF occupancy in the chromatin landscape of tTregs and Tconvs, provides a computational tool to infer the GRM governing human regulatory T cells with meaningful biology significance.

The AP-1 GRM is defined by its higher differential binding to upregulated genes in tTregs. AP-1 plays a central role in T-cell activation (Yukawa et al. 2019), Th differentiation, and T-cell anergy (Samstein et al. 2012). In murine Tregs it may promote Foxp3 expression through binding to its regulatory sites (Ogawa et al. 2014; Samstein et al. 2012). The AP-1 GRM includes BATF and the repressor BACH2. In mice, BACH2 interacts with AP-1 members at the shared consensus sites in thymic-derived Tregs (Sidwell et al. 2020), and targets lineage super-enhancers (Roychoudhuri et al. 2013). Our data show that BACH2 is downregulated in tTregs, so its repressive function is likely alleviated in tTregs. Thus, our data support a role for the AP-1 family in the establishment of human tTreg cells (Trujillo-Ochoa et al. 2023).

The KLF/SP GRMs may represent a more diverse mechanism in differential binding. KLF factors regulate multiple aspects of T-cell and lymphocyte biology, such as development, differentiation, trafficking, maturation, and quiescence (Hart et al. 2012). Consistently, our data indicates direct upregulation of diverse members of the NF-κB pathway by the KLF/SP cluster. We speculate that KLF/SP combine TF expression levels with alternating differential binding to refine derepression/activation of specific transcripts in the tTreg gene signature. Eg, whilst KLF6 overexpression drives the upregulation of its bound targets (McConnell and Yang 2010), it may be out-competed in binding to downregulated targets by less expressed repressors, *e.g.*, KLF9, KLF11, or SP3. Of note, KLF6 role in lymphocyte biology remains unclear.

Finally, we uncovered a GRM resulting from ETS family TFs and down DEGs clusters. In mice, it has been suggested that Foxp3 exploits the enhancer landscape bound by Ets factors to specify the Treg lineage (Samstein et al. 2012). In addition, Elf4 facilitates thymic Foxp3 expression (Mouly et al. 2010), which is consistent with ELF4 expression and binding in human tTregs.

We believe that our original strategy to focus on mature thymic regulatory and conventional cells represents an advantage for the identification of the gene modules governing human Tregs. Our approach limits confounding factors generated by activated / differentiated cells in the periphery, since human conventional CD4 T cells may express Treg markers upon TCR stimulation, challenging Treg lineage isolation (Caramalho et al. 2015a). Moreover, a significant component of peripheral Tregs may be regulatory T cells induced from conventional CD4 T cells, which are thought to be more plastic and to acquire more easily conventional profiles according to environmental conditions (Silva and Sousa 2016; Silva et al. 2016; Kanamori et al. 2016). Additionally, thymic Tregs are known to be particularly relevant in the control of autoimmunity, in part due to their enrichment in self-reactive TCRs (Caramalho et al. 2015a; Lee et al. 2012). On the other hand, a limitation of our study is the low number of replicates used to generate the GRM.

The omnigenic model (Boyle et al. 2017) proposes that genomic variants with small molecular effects may contribute toward a complex trait. Their cumulative impact would be relayed to a set of core genes through cell-specific regulatory networks. We showed that GRM mapped to mutational hotspots in healthy subjects and were significantly enriched in rare variants in CVID patients. These findings are strongly suggestive of the biological relevance underlying the TF-target interactions they codify. It is therefore reasonable to attempt the stratification of CVID patients based on GRM variant enrichment. We found an immediate classifier in AP1_upDEG, which is consistent with the recognised role of this protein complex in Treg development (Ogawa et al. 2014; Trujillo-Ochoa et al. 2023) and the upregulation of the transcription factor *BATF* in these cells. This is followed by KLF GRM or ETS_downDEG GRM, defining a hierarchy for patient segregation. However, at the time of this study we could not observe a direct correlation between these clusters and clinical manifestations in the respective patients, which may be in part related to the overlap of clinical phenotypes in CVID patients with immune dysregulation (Motta-Raymundo et al. 2022). Another confounding factor is that the disease is likely progressive and some patients might develop different complications overtime (Janssen et al. 2021). Future studies should validate our data and further investigate the variant landscape of other clinical contexts.

Taken together, these results suggest an application for GRM in the prioritisation of rare variants, and a possible alternative to expression levels as a function/impact classifier. Moreover, accumulation of SNVs associated to specific sets of GRM genes could be used to infer candidate pathways to be further explored to disentangle the polygenic basis of complex disorders and identify potential targets for personalised therapy.

Here, we demonstrated how analysing differential binding information extracted from bulk ATAC-seq from contrasting cellular types is a valuable strategy to uncover gene regulatory modules. We generated a resource of key gene regulatory modules governing the human Treg expression through their signature chromatin accessibility in the thymus. We found the application to the CVID genomic context to be particularly suitable whilst probing the GRM strategy in prioritising variants in complex immune diseases. The results support a broader application of the GRM model to other complex disorders, and to unlock the potential of whole-genome sequencing, namely by helping to evaluate variants of uncertain significance and/or their combined impact at an individual level.

## METHODS

Our aim was to uncover the gene regulatory modules (GRM) underlying the human thymic Treg signature and demonstrate their potential use in prioritising rare variants in diseases with a main polygenic basis like CVID. Briefly, we generated chromatin landscape and expression profiles of CD4 single-positive (CD4SP) Treg and Tconv cells (“tTregs” and “tTconvs”) purified from the human thymus from healthy donors. The GRM model was based on the clustering of Transcription Factor-Differentially Expressed Gene (TF-DEG) pairs via TF differential binding between lineages at accessible chromatin regions. We then assessed the potential biological relevance of GRM genes by their enrichment in rare variants found in CVID patients. Below the description of our methods, additional information may be found in Supplementary Materials.

### Human samples

Thymic samples were obtained during paediatric reconstructive cardiac surgery, using tissue that would be otherwise discarded (3 male and 3 female children, between 1 and 27 months of age, without evidence of immunodeficiency or syndromic diseases). Peripheral blood from 35 patients with a clinical diagnosis of CVID (Bonilla et al. 2016; Tangye et al. 2022), under follow-up at the adult PID outpatient clinic of Centro Hospitalar Universitário Lisboa Norte (CHULN)/Hospital de Santa Maria were selected based on their severe inflammatory/ autoimmune clinical phenotypes, as depicted in Supplemental Table S6. All participants or their legal representatives provided written informed consents. The study was approved by the ethical board of CHULN/Faculdade de Medicina da Universidade de Lisboa (FMUL)/Centro Académico Medicina de Lisboa (CAML).

### Cell Sorting and Flow Cytometry Analysis

Thymocytes isolated by Ficoll-Hypaque (GE Healthcare) from cell suspensions obtained by thymic tissue manual dispersion, were sort-purified to obtain mature CD4 single-positive (CD4SP) regulatory (Tregs) and conventional (Tconvs) thymocytes (purities above 95%), based on the surface expression of CD4, CD8, CD27, CD25 and CD127 using a FACS Aria III (BD Biosciences), as illustrated in Fig. S1A. CD3 was intentionally not used to avoid possible signalling, but the sorting strategy was validated in parallel using CD3 and intracellular Foxp3 in a Fortessa flow cytometer (BD Biosciences) using staining protocols previously described (Nunes-Cabaço et al. 2010), and the antibodies listed in Supplementary Materials. Analysis was performed using FlowJo v10 software.

### RNA-seq and differential expression analysis

RNA was extracted from cell pellets of 600,000 sorted tTregs and tTconvs from three different thymuses, using the AllPrep DNA/RNA kit (QIAGEN) and following the manufacturer’s instructions. Libraries were built selecting for polyadenylated RNA after depleting ribosomal fraction and then sequenced at both ends by high-throughput parallel sequencing (RNA-seq) in an Illumina Hiseq4000 sequencer (BGI Tech Solutions, Hong Kong, China). Raw sequencing was processed and analysed with SAMtools (Danecek et al. 2021), and sequence quality assessed with FastQC (see Table S1 in Supplementary Materials). The resulting ca. 200 million paired-end reads per biological replicate (PE100) were uniquely mapped and annotated to the human genome (hg38) with TopHat2 (Kim et al. 2013) and transcript expression quantified with R package HTSeq (Anders et al. 2015) (Count Per Million, CPM), with exclusion of genes with less than 1 CPM in more than 2 libraries. Libraries were scaled by Trimmed Mean of M-values (TMM) normalisation and corrected for heterogeneity of samples specific to contrast matrix with weighted scaling based on voom (Ritchie et al. 2015), followed by the quantification of Differential Expression between tTregs and tTconvs with R package edgeR (Robinson et al. 2010). Finally, we fitted multiple linear models by lmFit. Conversion between annotations was made with R biomaRt (Durinck et al. 2009). Differential Gene Expression threshold set between tTregs and tTconvs at log_2_FC > ±2, with FDR < 0.05 (Supplemental Table S1).

### ATAC-seq libraries, Regions of Open Chromatin (ROC) and Differential Chromatin Accessibility (DCA)

ATAC-seq was performed following the Omni-ATAC protocol *(Corces et al. 2017)* with minor modifications, using 5×10^4^ sorted tTreg or tTconv cells purified from 3 different thymuses. Cells were lysed for 3 minutes on ice, in 50uL of ATAC-Resuspension Buffer (10mM Tris-HCl pH 7.4, 10mM NaCl, 3mM MgCl_2_) containing 0.1% NP40, 0.1% Tween-20, and 0.01% Digitonin. tn5 tagmentation was performed using TDE1 Enzyme and Buffer TD (Illumina) at 37°C for 30 minutes, shaking at 1000rpm. After purification with a MinElute PCR Purification Kit (Qiagen), samples were amplified with NEBNext High Fidelity 2x PCR Master Mix (New England Biolabs) (Buenrostro et al. 2015). Final PCR reaction was then purified with a MinElute PCR Purification Kit followed by size-selection (150bp-1000bp using Ampure XP beads, Beckman Coulter). Sequencing was performed using a MGISEQ-2000 (BGI Tech Solutions), yielding a total sequencing depth between ∼200 and 600 million PE50 reads, and sequencing quality was assessed using FastQC. Reads were uniquely mapped to hg38 using Bowtie2 (Langmead and Salzberg 2012) and adapted for peak calling by MACS2 (Zhang et al. 2008) using inhouse pipeline, namely by converting to appropriate formats and correcting tn5 shift. Peaks from all samples were merged to create the total landscape of Regions of Open Chromatin (ROC) and signal assigned with BAMscale (Pongor et al. 2020). These peaks were annotated to Nearest Transcription Start Site with PeakAnalyzer (Salmon-Divon et al. 2010), using GTF annotation for hg38. To determine chromatin accessibility and its variation between tTregs and tTconvs (Differential Chromatin Accessibility, DCA), we applied the same tools, method, normalisations/rescaling as those described above for RNA-seq libraries, with the Peak_ID of each Region of Open Chromatin serving as the anchor for signal computation (Supplemental Table S2). DCA vs DEG linear regression analysis calculated with MM-type estimators (“lmrob” function of robustbase R package) to correct for data heteroscedasticity.

### Digital Genomic Footprinting and Transcription Factor Binding analysis

We used the TOBIAS framework 0.12.6 (Bentsen et al. 2020) to quantify protein occupancy in Regions of Open Chromatin (ROC, Supplemental Table S2), “treg_score” and “tconv_score” and then identify the underlying consensus motif (“motif_score”, which measures the sequence match) at each of the genomic footprints - or TF binding sites (TFBS). Continuous footprint scores with *p* < 0.01 across accessible chromatin regions were considered ‘bound’ by a transcription factor. Transcription factor motifs within ROC were identified using the Positional Weight Matrixes (PWMs) in the JASPAR Core database (Fornes et al. 2020; Khan et al. 2018). We selected 639 motif profiles matching “Homo Sapiens species” + “Latest Version”. Hypergeometric testing of cell-specific segments with TFBS were performed against 1,000 random equal-size sampling of the universe of human thymic TFBS, assuming a normal distribution. Supplemental Table S3 lists TFBS data. Further *in silico* epigenomics analysis described in Supplementary Materials.

### Differential Binding Cluster analysis and Gene Regulatory Modules

To assess the existence of patterns between the TFBS and the tTreg Signature genes a matrix was built with the genes as rows and TFBS as columns. After scaling the matrix by rows and calculating the optimal number of clusters through elbow and silhouette methods, two *k*-means algorithms were run simultaneously, one for the rows, one for the columns. Differential binding densities were calculated as the sum of all mean differential binding quantified in each Gene Regulatory Module divided by its surface area (number of DEGs * number of TFs), and the top 6 were selected (excluding the single-TF CTCF module).

### Whole-genome sequencing (WGS) and Variant calling

Genomic DNA extracted from peripheral blood of 35 CVID patients and sequenced to an average read depth of 30x (BGI-Shenzhen). The sequence reads were mapped to the reference GRCh37 genome using the BWA-MEM aligner, version 0.7.17 (Li 2013). Downstream processing was performed with SAMtools, and Picard Tools (http://broadinstitute.github.io/picard). Additionally, we used WGS data from 35 “Healthy Control” (HC) GRCh38 genomes, gender-balanced (17M/18F) and randomly selected amongst the subset “Iberian populations in Spain”, download from the International Genome Sample Resource (IGSR) (Clarke et al. 2017), generated from blood cells of healthy individuals. GATK4 germline short variant calling pipeline (Poplin et al. 2017), following Best Practices, VCFanno (Pedersen et al. 2016) and Ensembl Variant Effect Predictor (McLaren et al. 2016), and genome Aggregation Database (Karczewski et al. 2020) were used for haplotype calling, filtering, and annotation of single-nucleotide variants (SNV) found at the loci of genes expressed in tTreg and tTconvs and at regions of open chromatin associated with these genes. Calls with a read coverage of <30x were filtered out. Synonymous variants in gene loci were excluded and the remainder were only included for allelic frequency in non-Finnish Europeans (AF_NFE) < 0.01. The pipeline was adapted from Motta-Raymundo et al (Motta-Raymundo et al. 2022) (Supplementary Materials). Variants were analysed within the universe of genes expressed in our CD4SP RNAseq data (n=11,596) and with the subsets pertaining to the tTreg signature (n=1,357 DEG), or the different identified GRM (total n=368 DEG).

### Other data visualisation

Custom tracks were obtained by loading the respective RNA-seq and/or ATAC-seq bigwig files into IGV (Robinson et al. 2011). All heatmaps were created with the aid of the R “ComplexHeatmap” package. The other charts were created with the R packages “ggplot2”, or “enhancedVolcano”. Visual representations of the gene regulatory networks (cluster network graphs) were generated with Cytoscape v3.8.2 (Shannon et al. 2003) using the force-directed Compound Spring Embedder (Cose) layout followed by a removal of overlaps between the nodes (yFiles Remove Overlaps).

### Quantification and Statistical Analyses

All quantifications and statistical significance were calculated with R/Bioconductor, unless indicated otherwise. False-Discovery Rate, FDR, corresponds to adjusted p-value by multiple testing with Benjamin-Hochberg correction. The cut-off for expression of 2-fold change warrants the selection for differences with potential biological relevance and was overall on-pair with the chosen FDR<0.05.

## DATA ACCESS

Further information and requests for resources and reagents should be directed to and will be fulfilled by the corresponding author, Alexandre ASF Raposo. Human CD4SP T-cell ATAC-seq and RNA-seq raw and processed data generated in this study have been submitted to ArrayExpress. Accession numbers and reviewer logins (also listed in the Reagents and Tools table), respectively: E-MTAB-11220 and E-MTAB-11211. Publically available data used are listed in Materials and Tools table. Patient WGS raw data and corresponding VCF files (hg19) for the gene loci here described in CVID and healthy controls have been submitted to FEGA PT. All original relevant code has been deposited at GitHub (https://github.com/AESousaLabIMM/Kmeans_TOBIAS_CD4Thymus_paper). Links are listed in the Supplemental Materials.

## ACKNOWLEDGEMENTS

We thank Miguel Abecasis MD, Rui Anjos MD, and Hospital de Santa Cruz-Centro Hospitalar Lisboa Ocidental, and all parents and children for the access to thymic tissue; and the iMM flow cytometry facility; Miguel Ângelo-Dias, Zoe Junginger, Adriana Motta-Raymundo, and Diana Santos for technical assistance; and Bruno Silva-Santos, Luís Graça, Nuno Barbosa-Morais, Diogo S. Castro, and Saumya Kumar for the critical reading of the manuscript. This work was supported by grants PAC-PRECISE-LISBOA-01-0145-FEDER-016394 and LISBOA-01-0145-FEDER-007391, co-funded by FEDER through POR Lisboa 2020 - Programa Operacional Regional de Lisboa PORTUGAL 2020 and Fundação para a Ciência e a Tecnologia (FCT); PTDC/MED-IMU/0938/2020 to AES, funded by the Portuguese government through FCT. PR, AG-S, and HNC were co-funded by FCT and FEDER; SP was funded by GenomePT (PINFRA/22184/2016/POCI-01-0145-FEDER-022184). YT received an ENLIGHT-TEN PhD fellowship (MSCA-ITN-2015-675395 - H2020) from Marie Sklowdowska-Curie-COFUND Action; AASFR was funded by FCT (CEECIND/01474/2017).

## AUTHOR CONTRIBUTIONS

AASFR, PR, and AES designed the study; ARMA, AGS, HNC, PR, and YT collected data; AASFR, PR, SP, and MEB performed the computational analyses; SLS collected the clinical data; AASFR, PR, ARMA, and AES analysed and interpreted the results; AASFR and AES supervised the study; AASFR and AES wrote the manuscript. All authors discussed the results and contributed to the final manuscript.

