## Supplemental Material and Figures for "Decoding mutational hotspots in human disease through the gene modules governing thymic regulatory T cells"

**This PDF file includes:**

Supplemental Methods
Figures S1 to S7
Legends for Supplemental Tables S1 to S9
Supplemental Table S10
Supplemental Material References

**Other supporting materials for this manuscript include the following:**

Supplemental Tables S1 to S9

**Supplemental Methods**

**Human sample collection**

Blood samples were obtained from CVID patients. The diagnosis of Granulomatous Lymphocytic Interstitial Lung Disease (GLILD), Liver Regenerative Nodular Hyperplasia (LRNH), and Gastric Cancer, were based on organ biopsies. CVID-associated Enteropathy was defined as chronic inflammation in gut histology and/or malabsorption in the absence of pathogen isolation in stools or biopsies. Lymphoproliferation was defined by adenomegalies (lymph nodes larger than 1cm diameter in ≥2 lymphatic chains in clinical and/or imaging exams) and/or splenomegaly (longitudinal spleen diameter superior to 15 cm by computed tomography or ultrasonography). The standard clinical criteria were used for the diagnosis of organ autoimmunity and cytopenia.

**ATAC-seq libraries and data generation**

Samples amplification with index adapters from:
i5- AATGATACGGCGACCACCGAGATCTACACTCGTCGGCAGCGTCAGATGTG
i7- CAAGCAGAAGACGGCATACGAGATNNNNNNNNGTCTCGTGGGCTCGGAGATGT (barcodes identified as NNNNNNNN).

**Regions of Open Chromatin (ROCs) and Differential Chromatin Accessibility**

Sequencing data processing for peak calling using inhouse pipeline (can be provided upon request): removing duplicate and mitochondrial reads, selecting properly paired reads, sorting and indexing, converting BAM into BEDPE format, correcting tn5 shift.
Finally running MACS2 command with the following parameters:
*macs2 callpeak -t ${bam} -f BAMPE -g hs -q 0.05 --nomodel *
*--extsize 200 --shift -100 -n ${bam} --outdir PEAKS*
In parallel, reads were normalised for visualisation in Integrative Genomics Viewer (1) as pile-up BigWig custom tracks with BAMscale (2). These custom tracks can be downloaded from ArrayExpress.

**Super-enhancer, enhancer, and promoter regions**

In the absence of thymic chromatin data, we resorted to available peripheral Treg and Tconv human datasets to define Treg super-enhancer, enhancer, and promoter, obtained from the Roadmap Epigenomics project (CD4+CD25+CD127- Treg Primary Cells, and CD4+CD25- Th Primary Cells). Enhancers and promoters were obtained from corresponding chromatin states data available. To generate a map of super-enhancers, we stitched the enhancers (ENCODEENCSR577GVS and ENCSR546SDM, ENCFF794AWE and ENCFF058CQG, respectively) using ROSE (https://github.com/stjude/ROSE) and separated the result from typical enhancers using sequencing data in BAM format, given a GFF file of previously identified constituent enhancers. Stitching distance was set to regions within 12.5kb of one another, with a TSS exclusion distance of 4kb. Segments of super-enhancers, enhancers, and promoters specific to Treg or Tconv were obtained using bedtools intersect (https://bedtools.readthedocs.io/en/latest/content/bedtools-suite.html). The p-values of the overlap of TFBS with genomic features were calculated against 1,000 random equal-size sampling of the universe of human thymic TFBS, assuming a normal distribution for the successful tests.

**Other data sets**

Identities of genes associated to Primary Immunodeficiency and Common Variable Immunodeficiency were obtained from a compilation of 2022 updates to IUIS Phenotypical Classification for Human Inborn Errors of Immunity.

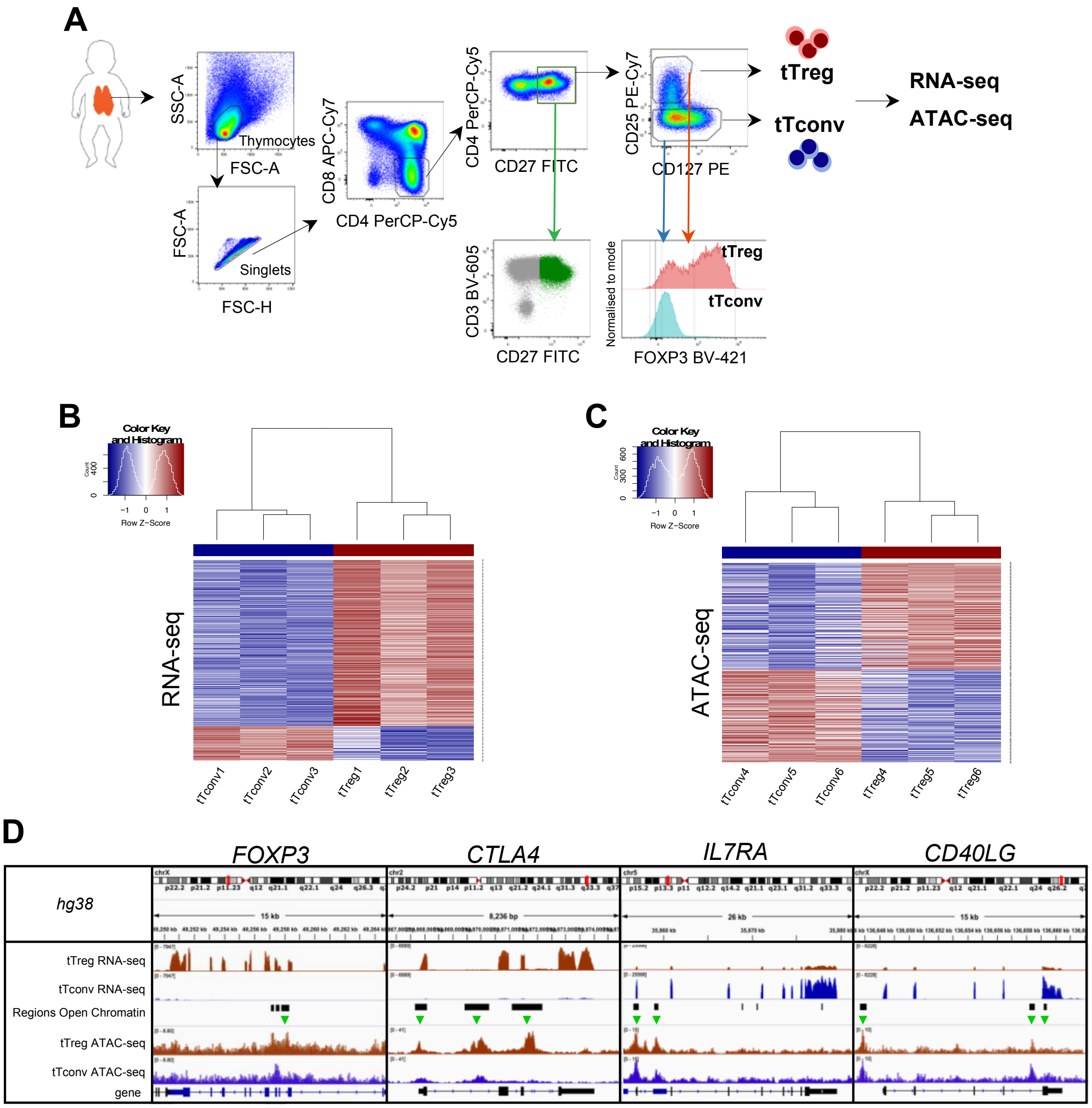


**Fig. S1. The human thymic Treg signature and its correlation with chromatin accessibility (related to Figure 1).** A. Validation of the sorting strategy showing CD3bright and CD27 expression in CD4SP and levels of *FOXP3* in the sorted tTregs and tTconvs. B-C. RNA-seq (B) and ATAC-seq (C) for top 1,000 genes and top 1,000 Regions of Open Chromatin (ROC), ranked by respective fold-change, segregating between tTregs (red sidebar) and tTconvs (blue sidebar). D. Profiles of raw expression and accessibility to chromatin at representative genes in tTreg (red) and tTconv (blue); Top track - chromosome localisation; Bottom track - Gene: black for sense; blue for antisense; Green arrows indicate relevant ROC with higher accessibility in tTregs, which are particularly striking for *FOXP3*, mainly within introns overlapping the Conserved Non-Coding Sequences (CNS) targeted by its regulators (refs 12,33), for *CTLA4*, at the promotor and two introns, for *CD40LG*, at both genic and upstream loci of the corresponding Transcription Start Site (TSS), and for the *IL7RA* promotor region.

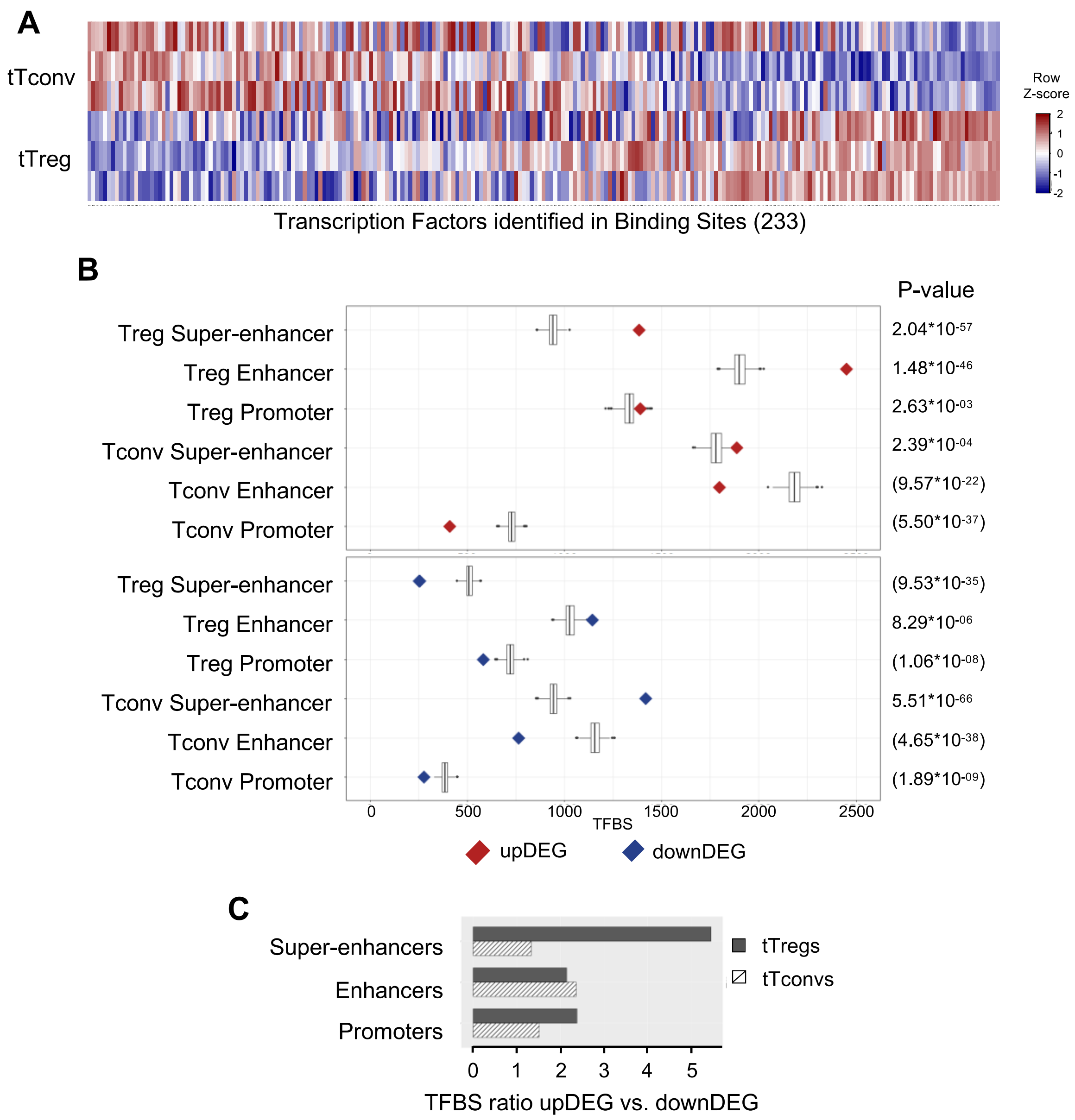


**Fig. S2. Treg-specific super-enhancers and enhancers, but not promoters, are enriched in tTreg TF binding sites (TFBSs), particularly when these sites are potentially activating transcription of tTreg signature up-regulated genes (related to Figure 1).** A. Gene expression in each of the three replicates in tTregs and tTconvs data sets for all transcription factors that can be identified through a position-weight matrix (PWM) in the JASPAR database (233 of 639, $log_2CPM$ shown in row Z-score). B. Frequencies of TF Binding Sites (TFBS) mapping to specific Tconv, or specific Treg super-enhancer, enhancer, and promoter regions; Red, TFBS associated to the 513 upDEG; Blue, TFBS associated to the 312 downDEG; Boxplots: distribution of 1,000 random same-size sampling of TFBS universe (binomial test); Right: corresponding p-values. C. TFBS mapping to Treg lineage-specific super-enhancers, enhancers, and promoters (from B.); Bar length: ratio in TFBS between upDEG and downDEG; dark grey, tTregs; hatched, tTconvs.

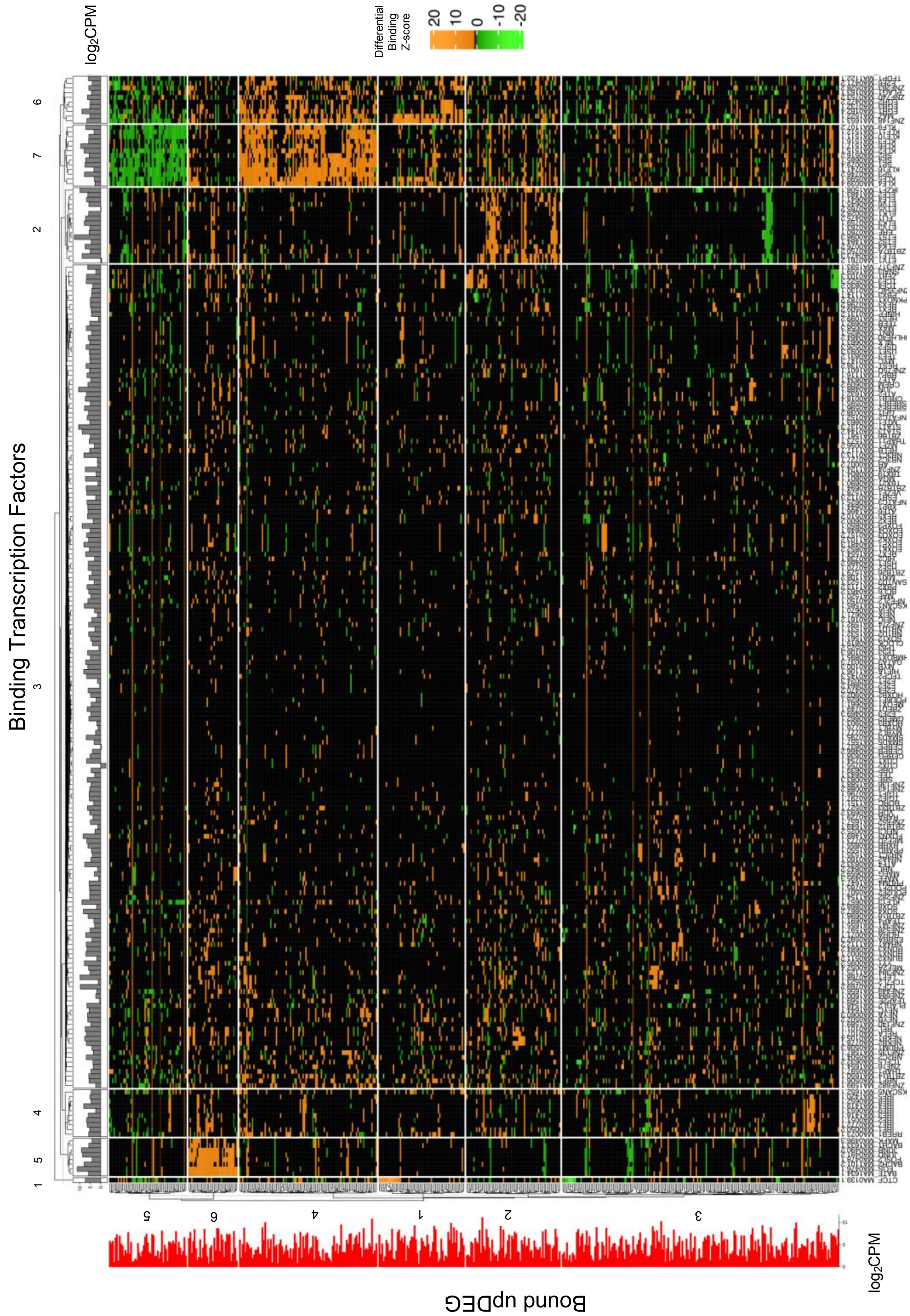


**Fig. S3. TF Differential Binding reveals main Gene Regulatory Modules controlling the tTreg Signature in upDEG (related to Figure 2).** Expanded version of heatmap shown in the top of Figure 2. Column sidebar: transcription factors expression levels in tTregs ($log_2$CPM, grey);

Row side bar: upDEG expression levels in tTregs ($log_2$CPM, red).

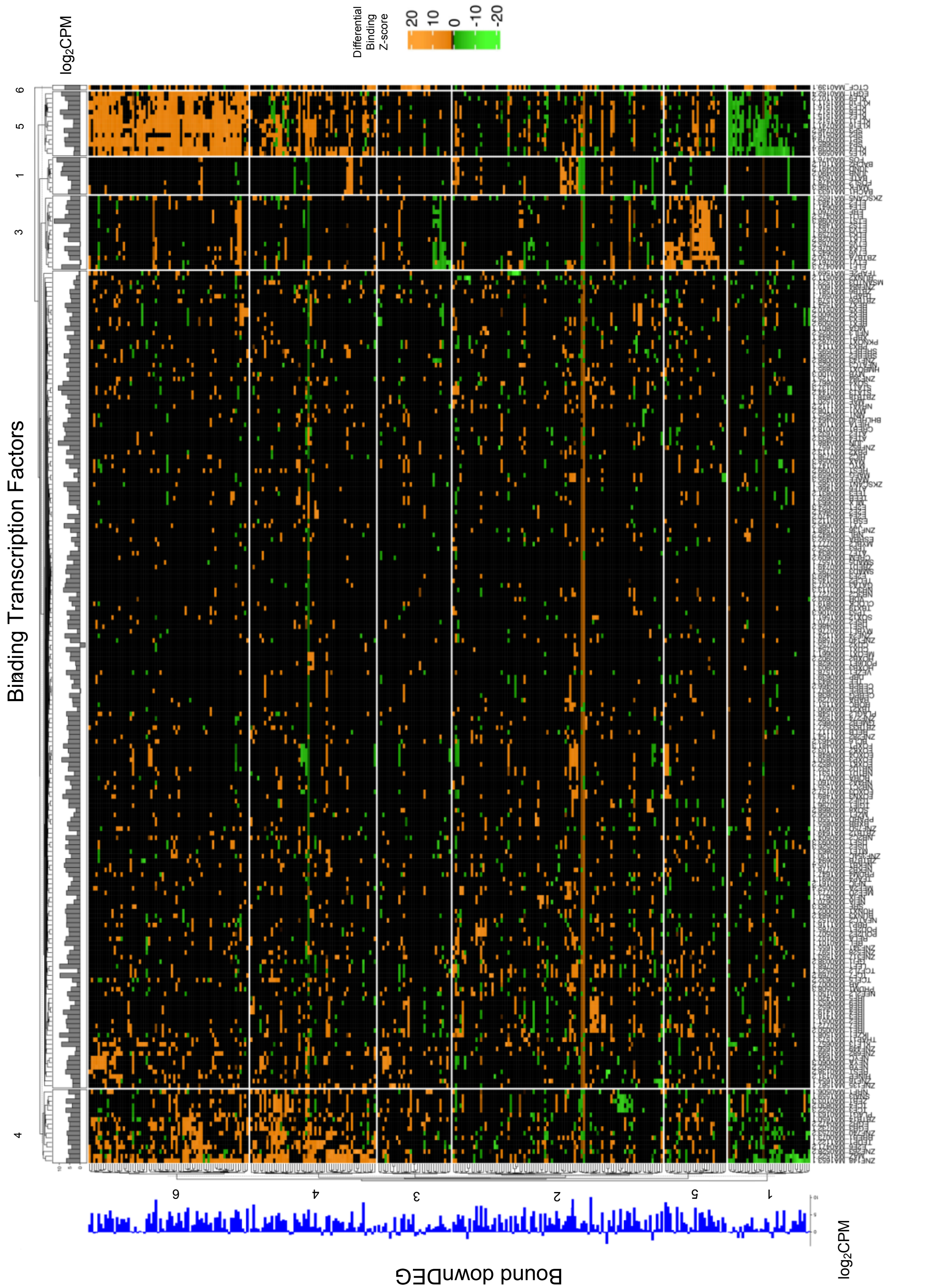


**Fig. S4. TF Differential Binding reveals main Gene Regulatory Modules controlling the tTreg Signature in downDEG (related to Figure 2).** Expanded version of heatmap shown in the bottom in Figure 2; Column sidebar: transcription factors expression levels in tTregs ($log_2$CPM, grey); Row side bar: downDEG expression levels in tTregs ($log_2$CPM, blue).

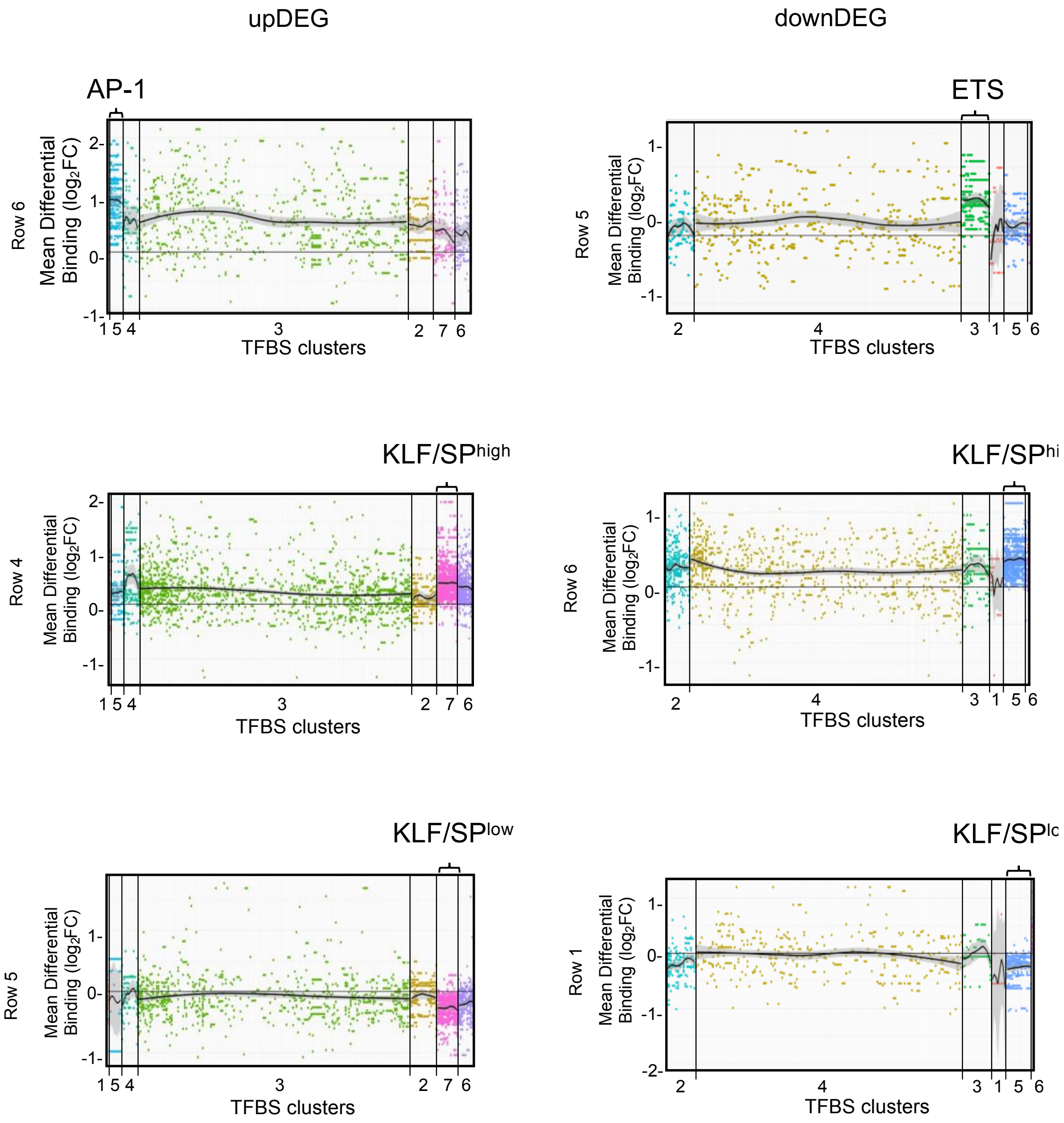


**Fig. S5. Significant Gene Regulatory Modules and respective Transcription Factor Differential Expression vs Transcription Factor Differential Binding to targets (related to Figure 2).** Mean differential binding by each transcription factor to each target in the significant Gene Regulatory Modules found (scatter plot); Transcription Factor Binding Site (TFBS) clusters are coloured differently for clarity, and asterisk indicates cluster with highest score for differential binding density (see Methods); Black line, LOESS over TFBS clusters (grey shade, Confidence Interval); Left, in upDEG, right, in downDEG.

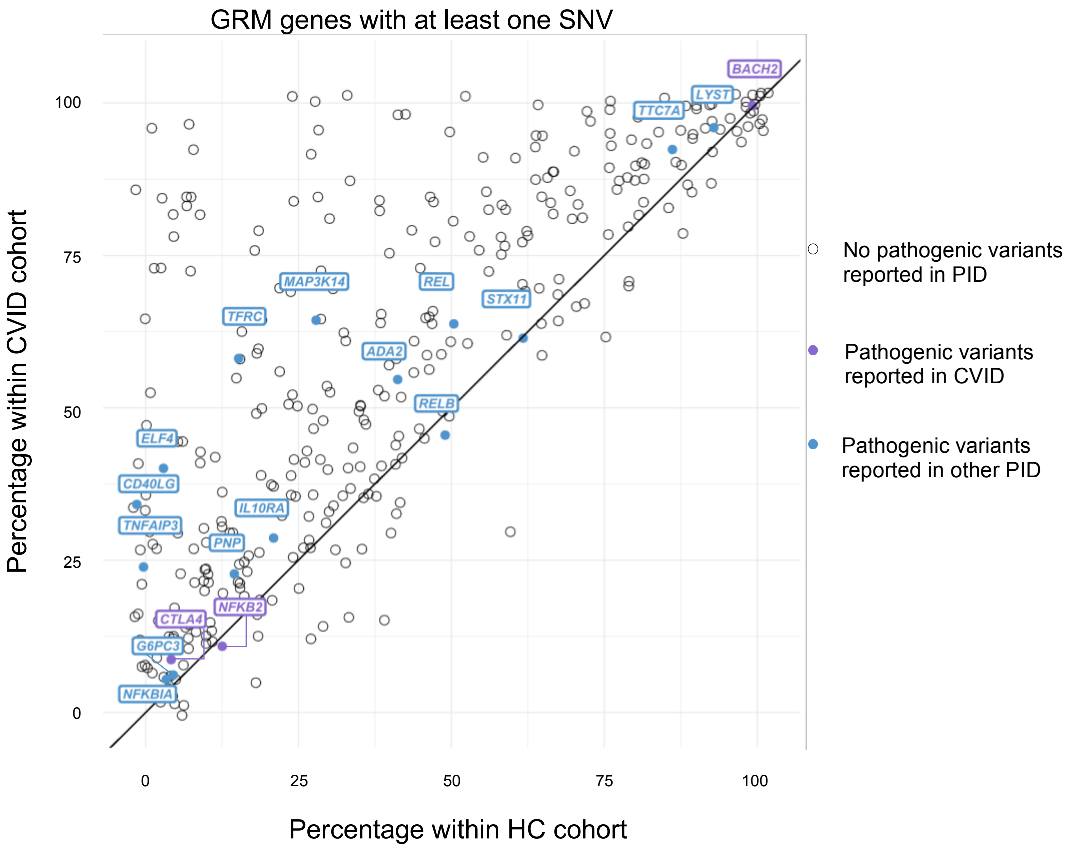


**Fig. S6. Analysis of the proportion of genes of the Gene Regulatory Modules (GRM) of thymic Treg (tTreg) with at least one SNV in the CVID versus healthy control cohorts (related to Figure 5)**. Proportion of individuals within the CVID and HC cohorts with at least one SNV for each GRM gene; Line indicates equal prevalence; Each dot represents a gene (N=355; genes with no SNV in both cohorts not shown), and those with previously reported pathogenic mutations associated to CVID are highlighted in purple and to other primary immunodeficiencies (PIDs) in blue (from The 2022 Update of IUIS Phenotypical Classification for Human Inborn Errors of Immunity, https://doi.org/10.1007/s10875-022-01352-z; Human Inborn Errors of Immunity: 2022 Update on the Classification from the International Union of Immunological Societies Expert Committee, https://doi.org/10.1007/s10875-022-01289-3).

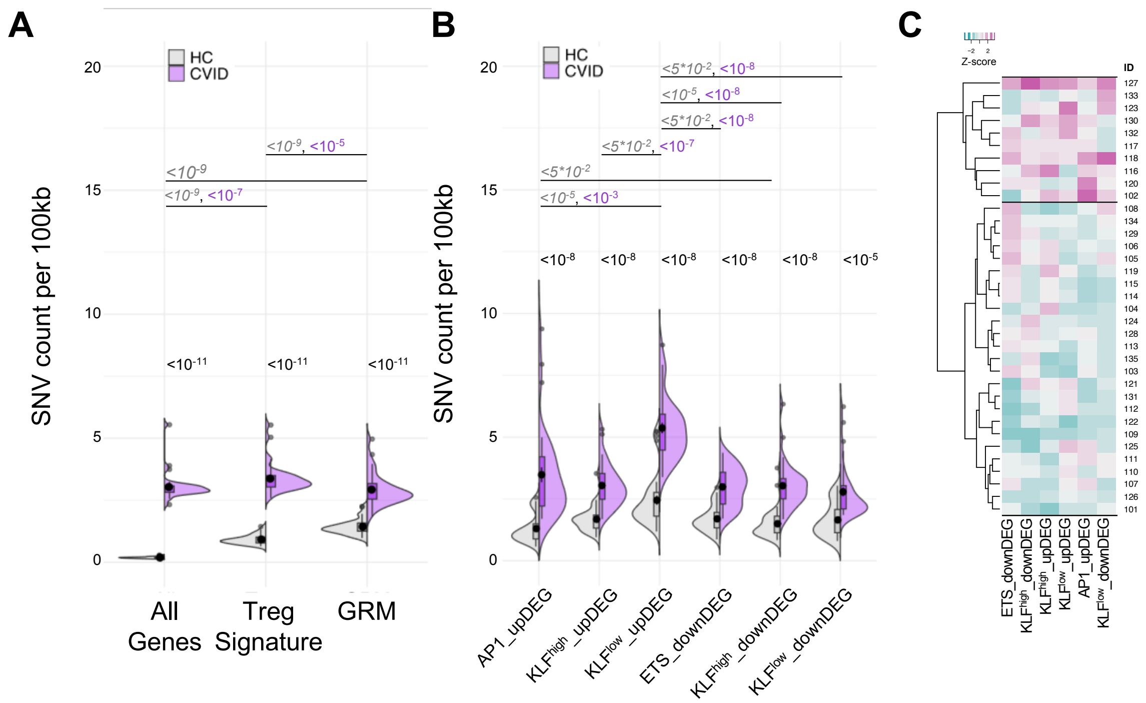


**Fig. S7.** ***The Gene Regulatory Modules (GRM) of thymic Treg (tTreg) are enriched in rare variants* (related to Figure 5)**. A-B. Comparison of mutation load or variant density (SNV per 100k) in gene loci between the healthy control cohort (HC, grey) and Common Variable Immunodeficiency cohort (CVID , purple): distribution in all genes expressed in tTregs and tTconvs; in genes of the tTreg Signature (DEG); and in genes forming the GRM are shown in the violin plots of (A); and in gene loci for each of the GRM in (B). C. Clustering of CVID patients by mutational load in each GRM; the two major clusters are mostly due to differences in AP1_upDEGs and KLF GRM (darker magenta, higher variant density; darker cyan, lower variant density).

**Supplemental Table S1 (separate file).** Thymic Treg Signature

**Supplemental Table S2 (separate file).** Regions of Open Chromatin associated to Differential Expressed Genes

**Supplemental Table S3 (separate file).** Transcription Factor Binding Sites found in tTregs

**Supplemental Table S4 (separate file).** FOXP3 direct targets amongst Differentially Expressed Genes and respective Transcription Factor Bindings Sites; Transcription Factor Binding Sites at other Differentially Expressed Genes.

**Supplemental Table S5 (separate file).** List of all clusters of Transcription Factors and Differentially Expressed Genes based on Differential Binding Analysis

**Supplemental Table S6 (separate file).** Clinical and Epidemiological data of patients with Common Variable Imunodeficiency (CVID)

**Supplemental Table S7 (separate file).** Fractions of genes with at least one SNV: analysis of all expression; Treg Signature; and GRMs in CVID and healthy individuals.

**Supplemental Table S8 (separate file).** SNV count per 100kb (density): analysis of all expression; Treg Signature; and GRMs in CVID and healthy individuals.

**Supplemental Table S9 (separate file).** Proportion of genes of the Gene Regulatory Modules (GRM) of thymic Treg (tTreg) with at least one SNV in the CVID or healthy control cohort.

**Supplemental Table S10. Materials and Tools**

| REAGENT or RESOURCE | SOURCE | IDENTIFIER |
|---|---|---|
| **Antibodies** | | |
| Alexa Fluor® 647 Mouse monoclonal anti-Bcl-6 | BD Biosciences | Cat# 561525; RRID: AB_10898007 |
| Mouse monoclonal APC anti-human CD198 (CCR8) | BioLegend | Cat# 360609; RRID: AB_2820017 |
| Mouse monoclonal anti-Human IL-7R alpha/CD127 PE-conjugated | R&D Systems | Cat# FAB306P-100; RRID: AB_2233759 |
| CD127 Monoclonal Antibody (eBioRDR5), eFluor 660, eBioscience™ | Thermo Fisher Scientific | Cat# 50-1278-42; RRID: AB_11217472 |
| Alexa Fluor(R) 700 mouse monoclonal anti-human CD127 (IL-7Ralpha) | Biolegend | Cat# 351344, RRID:AB_2566200 |
| Mouse monoclonal CD25 PE-Cy7 CE | BD Biosciences | Cat# 335824, RRID:AB_2868687 |
| Mouse monoclonal PE/Cyanine5 anti-human CD25 | Biolegend | Cat# 302608, RRID:AB_314278 |
| CD27 Mouse Monoclonal Antibody (O323), FITC, eBioscience™ | Thermo Fisher Scientific | Cat# 11-0279-42, RRID:AB_10669045 |
| Brilliant Violet 605(TM) anti-human CD3 antibody | BioLegend | Cat# 317322, RRID:AB_2561911 |
| CD39 Monoclonal Antibody (eBioA1 (A1)), PerCP-eFluor 710, eBioscience™ | Thermo Fisher Scientific | Cat# 46-0399-42, RRID:AB_10597271 |
| CD4 Monoclonal Antibody (RPA-T4), PerCP-Cyanine5.5, eBioscience™ | Thermo Fisher Scientific | Cat# 45-0049-42, RRID:AB_1518744 |
| Brilliant Violet 711(TM) anti-human CD4 antibody | BioLegend | Cat# 300558, RRID:AB_2564393 |
| BV510 Mouse Anti-Human CD45RA antibody | BD Biosciences | Cat# 563031, RRID:AB_2722499 |
| Brilliant Violet 785(TM) anti-human CD45RO antibody | BioLegend | Cat# 304234, RRID:AB_2563819 |
| PE/Dazzle(TM) 594 anti-human CD54 antibody | BioLegend | Cat# 353118, RRID:AB_2715946 |
| Mouse Anti-CD8 Monoclonal Antibody, APC-Cy7 Conjugated, Clone SK1 | BD Biosciences | Cat# 557834, RRID:AB_396892) |
| PE/Cyanine7 anti-human CD183 (CXCR3) antibody | BioLegend | Cat# 353720, RRID:AB_11219383 |
| PE/Dazzle(TM) 594 anti-human CD185 (CXCR5) antibody | BioLegend | Cat# 356928, RRID:AB_2563689 |
| FOXP3 Monoclonal Antibody (PCH101), eFluor 450, eBioscience™ | Thermo Fisher Scientific | Cat# 48-4776-42, RRID:AB_1834364 |
| Alexa Fluor(R) 700 anti-human/mouse/rat CD278 (ICOS) antibody | BioLegend | Cat# 313528, RRID:AB_2566126 |
| Mouse Anti-Human Ki-67 Antibody, Alexa Fluor® 647 Conjugated | BD Biosciences | Cat# 558615, RRID:AB_647130 |
| T-bet Monoclonal Antibody (eBio4B10 (4B10)), PE-Cyanine7, eBioscience™ | Thermo Fisher Scientific | Cat# 25-5825-82, RRID:AB_11042699 |
| TCR alpha/beta Monoclonal Antibody (IP26), APC, eBioscience™ | Thermo Fisher Scientific | Cat# 17-9986-42, RRID:AB_10597896 |
| Biological samples | | |
| Human Infant Thymus Sample | Hospital de Sta Cruz, Carnaxide, Portugal | T274 |
| Human Infant Thymus Sample | Hospital de Sta Cruz, Carnaxide, Portugal | T276 |

| | | |
|---|---|---|
| Human Infant Thymus Sample | Hospital de Sta Cruz, Carnaxide, Portugal | T277 |
| Human Infant Thymus Sample | Hospital de Sta Cruz, Carnaxide, Portugal | T349 |
| Human Infant Thymus Sample | Hospital de Sta Cruz, Carnaxide, Portugal | T350 |
| Human Infant Thymus Sample | Hospital de Sta Cruz, Carnaxide, Portugal | T353 |
| **Chemicals, peptides, and recombinant proteins** | | |
| Ficoll-Hypaque | GE Healthcare | Cat# 17544202 |
| Ampure XP beads | Beckman Coulter | A63880 |
| Critical commercial assays | | |
| AllPrep DNA/RNA kit | QIAGEN | Cat. No. / ID: 80284 |
| MinElute PCR Purification Kit | QIAGEN | Cat. No. / ID: 28004 |
| NEBNext High Fidelity 2x PCR Master Mix | New England Biolabs | M0541S |
| TDE1 Enzyme and Buffer TD kit | Illumina | 20034197 |
| **Deposited data** | | |
| Human reference genome NCBI build 37, GRCh37 | Genome Reference Consortium | https://www.ncbi.nlm.nih.gov/grc/human |
| Human reference genome NCBI build 38, GRCh38 | Genome Reference Consortium | https://www.ncbi.nlm.nih.gov/grc/human |
| gnomAD | Broad Institute Genome Aggregation Database | https://gnomad.broadinstitute.org/ |
| Iberian populations in Spain | International Genome Sample Resource[46] | https://www.internationalgenome.org/data-portal/population/IBS |
| ChIP-Seq analysis of H3K27ac in human CD25+ CD127- Treg cells_Jun-28-2012_38865 | Roadmap Epigenomics Project | https://www.encodeproject.org/experiments/ENCSR577GVS/ |
| ChIP-Seq analysis of H3K27ac in human CD25- Th cells_Jun-28-2012_29442 | Roadmap Epigenomics Project | https://www.encodeproject.org/experiments/ENCSR546SDM/ |
| FOXP3+ CD4SP Treg ATAC-seq | ArrayExpress | E-MTAB-11220 |
| FOXP3- CD4SP Tconv ATAC-seq | ArrayExpress | E-MTAB-11220 |
| FOXP3+ CD4SP Treg RNA-seq | ArrayExpress | E-MTAB-11211 |
| FOXP3- CD4SP Tconv RNA-seq | ArrayExpress | E-MTAB-11211 |
| **Software and algorithms** | (numbers correspond to references in main text) | |
| Bowtie2 | (3) | http://bowtie-bio.sourceforge.net/bowtie2/index.shtml |
| FlowJo v10 | | https://www.flowjo.com/solutions/flowjo/downloads |
| BWA-MEM | (4) | https://github.com/lh3/bwa |
| Samtools | (5) | http://samtools.sourceforge.net/ |

| | | |
|---|---|---|
| TopHat | (6) | https://ccb.jhu.edu/software/tophat/index.shtml |
| PeakAnalyzer | (7) | http://www.bioinformatics.org/peakanalyzer |
| Integrative Genomics Viewer | (1) | https://software.broadinstitute.org/software/igv/download |
| MACS2 | (8) | https://macs3-project.github.io/MACS/ |
| FastQC | | https://www.bioinformatics.babraham.ac.uk/projects/fastqc/ |
| R | R Foundation for Statistical Computing, Vienna, Austria. | https://www.r-project.org/ |
| robustbase | | https://robustbase.r-forge.r-project.org/ |
| HTSeqtools | (9) | https://doi.org/doi:10.18129/B9.bioc.htSeqTools |
| edgeR | (10) | https://doi.org/doi:10.18129/B9.bioc.edgeR |
| limma | (11) | https://doi.org/doi:10.18129/B9.bioc.limma |
| bioMart | (12) | https://doi.org/doi:10.18129/B9.bioc.biomaRt |
| Rank Ordering of Super-Enhancers (ROSE) | | https://github.com/stjude/ROSE |
| bedtools | | https://bedtools.readthedocs.io/en/latest/ |
| Tobias | (13) | https://github.com/loosolab/TOBIAS |
| ComplexHeatmap | | https://bioconductor.org/ |
| ggplot2 | | https://ggplot2.tidyverse.org/ |
| EnhancedVolcano | | https://doi.org/doi:10.18129/B9.bioc.EnhancedVolcano |
| Cytoscape v3.8.2 | (14) | https://cytoscape.org |
| BAMscale | (2) | |
| Kmeans_TOBIAS_CD4Thymus_paper | This study | https://github.com/AESousaLabIMM/Kmeans_TOBIAS_CD4Thymus_paper |
| Picard tools | | (9) |

| | | |
|---|---|---|
| GATK pipeline | (15, 16) | https://gatk.broadinstitute.org/hc/en-us/articles/360036194592-Getting-started-with-GATK4 |
| VEP | (17) | http://www.ensembl.org/info/docs/tools/vep/script/vep_download.html |
| VCFanno | (18) | https://github.com/brentp/vcfanno |
| **Other** | | |
| Hiseq 4000 | Illumina | |
| MGISEQ-2000 / DNBSEQ-G400 FAST | BGI Tech Solutions | |
| FACS Aria III | BD Biosciences | |
| LSRFortessa Cell Analyzer | BD Biosciences | |

14. Cytoscape: A Software Environment for Integrated Models of Biomolecular Interaction Networks (July 24, 2023).

15. Genomics in the Cloud [Book] (July 24, 2023).

16. R. Poplin, *et al.*, Scaling accurate genetic variant discovery to tens of thousands of sample (2017) https:/doi.org/10.1101/201178 (July 24, 2023).

17. W. McLaren, *et al.*, The Ensembl Variant Effect Predictor. *Genome Biology* **17**, 122 (2016).

18. Vcfanno: fast, flexible annotation of genetic variants | Genome Biology | Full Text (July 24, 2023).
